# Live-imaging reveals Coordinated Cell Migration and Cardiac Fate Determination during Mammalian Gastrulation

**DOI:** 10.1101/2023.12.19.572445

**Authors:** Shayma Abukar, Peter A. Embacher, Alessandro Ciccarelli, Sunita Varsani-Brown, Isabel G.W. North, Jamie A. Dean, James Briscoe, Kenzo Ivanovitch

## Abstract

Heart development involves the specification of cardiac progenitors at distinct stages and locations. Using live-imaging of mouse embryos between gastrulation and heart tube formation, we tracked individual mesodermal cells and reconstructed their lineage trees for up to five cell divisions. We found independent unipotent progenitors emerging at specific times, contributing exclusively to either left ventricle/atrioventricular canal (LV/AVC) or atrial myocytes. LV/AVC progenitors differentiated early to form the cardiac crescent, while atrial progenitors later generated the heart tube’s inflow tract during morphogenesis. We also identified short-lived bipotent progenitors with broad potential, illustrating early developmental plasticity. Sister cells from bipotent progenitors displayed greater dispersion and more diverse migratory trajectories within the anterior mesoderm than those from unipotent progenitors. Bipotent progenitors contributing to extraembryonic mesoderm (ExEm) exhibited the fastest and most dispersed migrations, whereas those giving rise to endocardial, LV/AVC, and pericardial cells showed a more gradual divergence, with late-stage behavioural shifts: endocardial cells increased in speed, while pericardial cells slowed relative to LV/AVC cells. Together the data reveal the regulation of individual cell directionality and cardiac fate allocation within the seemingly unorganised migratory pattern of mesoderm cells.

## Introduction

How cell fate specification and morphogenesis are coordinated in time and space to generate tissues and organs of unique forms and functions is central to developmental biology. This phenomenon is evident during gastrulation when mesodermal cells acquire diverse cardiac fates and engage in complex cell movements to generate spatial patterns, including a cohesive cardiac crescent, which transforms into a primitive heart tube.

In the gastrulating mouse embryo, tracking the derivatives of single progenitors by clonal analysis led to the finding that progenitors are assigned to specific anatomical locations in the heart prior to the formation of the heart fields ^1,2, 3, 4^. Unipotent *Mesp1+* progenitors are solely destined for the left ventricle (LV) and atria myocardium can be distinguished from unipotent endocardium progenitors ^4^. Additional clonal analysis of *Hand1+* progenitors located at the embryonic/extraembryonic boundary in the early gastrulating embryo identified bipotent and tripotent progenitors. These progenitors generated LV/AVC myocytes in addition to pericardium, epicardium, and extraembryonic tissues ^1^.

One limitation of clonal analysis, however, is that the history of the cells is *deduced* by analysing descendants at the endpoint. It does not allow the identification of the progenitors’ initial locations or subsequent migratory paths in the embryo. Single-cell tracking in live-imaging is needed for this and is the most rigorous approach to reconstituting cell lineages, identifying when cardiac progenitors become lineage-restricted during gastrulation and enabling migration analysis ^5^.

A recent live-imaging analysis uncovered the dynamics of mesodermal cell migration during mouse gastrulation ^6^. This analysis revealed that cells dispersed extensively in the embryo, with clearly separate movements of daughter cells, suggesting cell identity may not be fixed but instead influenced by the position of the cells at the end of the migration period. However, as Dominguez et al. discussed, the motility of mesodermal cells is unlikely to be completely random ^6^. There may be some regulation of directionality of individual cell migration to ensure progeny migrate to their correct locations and establish spatial patterns, including the cardiac crescent and distinct LV/AVC and atrial progenitor domains^7–9^. Indeed, a previous migration analysis showed mesodermal cell migrate with directionality during mouse gastrulation ^10^. Thus, early mammalian mesoderm migration may exhibit some degree of determinism. This phenomenon is reminiscent of an evolutionarily distinct species, the ascidian, in which a small number of genealogically related and determined heart progenitors migrate with predetermined directionally ^11^. However, it is not known whether progenitors adopt more stereotypical migratory trajectories, once committed to specific cardiac fates in the context of mammalian gastrulation.

Here, through long-term live imaging and single-cell tracking in mouse embryos, spanning from the initiation of gastrulation to the stages of heart tube formation, our goal was to reconstitute the lineage tree of cells and assess how the migratory paths of cells relate to their eventual cardiac fate within the seemingly unorganised migration pattern.

## Results

### Development and characterisation of *cTnnT-2a-eGFP* mice

To track cardiomyocytes in vivo, we developed a knock-in mouse reporter line *cTnnT-2a-eGFP* where the *eGFP* sequence is inserted downstream of the endogenous *cardiac troponin T* (*cTnnT*) loci. A virus-derived 2a self-cleaving peptide inserted between the *cTnnT* and *eGFP* coding sequence ensures co-expression of both cTnnT and eGFP proteins (Figure 1A)^12^. The *cTnnT-2a-eGFP* line was maintained as homozygotes. Animals are viable and indistinguishable from heterozygotes. Whole-mount immunostaining for cTnnT confirmed specific eGFP expression in cTnnT+ cardiomyocytes at E8 -heart tube stage- and E12.5 (Figure 1B-C).

**Figure 1.**
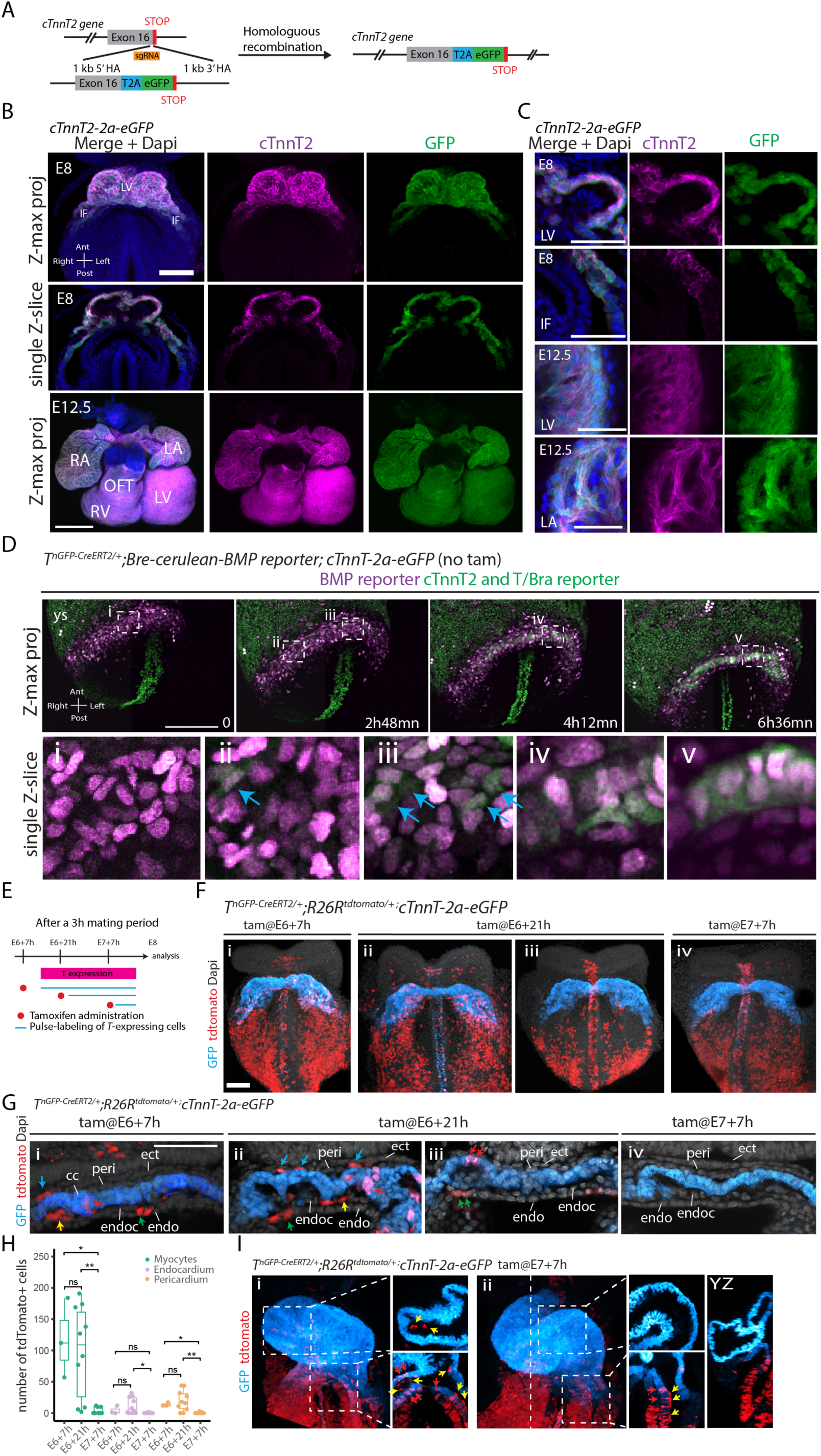
Characterization of the cTnnT-2a-GFP Line and Lineage Tracing of Early Cardiac Mesoderm. (A) CRISPR-Cas9 strategy to insert a 2a-eGFP cassette into the cTnnT2 gene. (**B-C**) Immunofluorescence for cTnnT2 in *cTnnT-2a-eGFP* hearts at E8 (scale: 100 μm) and E12.5 (scale: 200 μm). (**D**) Time-lapse sequence of a *T^nGPF-CreERT^*^2^*^/+^; Bre-cerulean; cTnnT-2a-eGFP* embryo (scale: 100 μm). Blue arrows points to *cTnnT-2a-eGFP+* cells. (**E**) Tamoxifen administration strategy. (**F**) *T^nGPF-CreERT^*^2^*^/+^;R26R^tdTomato/+^; cTnnT-2a-eGFP* embryos labeled at E6+7h, E6+21h, and E7+7h, analyzed at E8. Scale: 100 μm. (**G**) Single optical sections from (F) Scale: 100 μm. (**H**) Quantification of tdTomato+ cells (myocardial, endocardial, pericardial) in embryos labeled at E6+7h (n=3), E6+21h (n=10), and E7+7h (n=8). Data shown as individual embryos, analyzed with a Mann-Whitney U test. (**I**) Embryos labeled at E7+7h, analyzed at E8.5. IF: inflows, HT: heart tube, OFT: outflow tract, LV: left ventricle, RV: right ventricle, CC: cardiac crescent, peri: pericardium, ect: ectoderm, endoc: endocardium, endo: endoderm, YS: yolk sac. Arrows in (G) and (I) indicate cell types: blue for pericardium, yellow for endocardium, green for endoderm, and red for myocytes.

We first analyzed cardiac differentiation dynamics in real time using multiphoton live-imaging and the cardiomyocyte *cTnnT-2a-eGPF* reporter line (Figure 1D). We combined the *cTnnT-2a-eGPF* reporter with the *Bre:H2BCerulean* BMP reporter ^7^, which expressed cerulean in the lateral plate mesoderm. We found initial GFP positive cells appeared within the Bre-Cerulean positive lateral plate mesoderm at E7.5 - consistent with initial sparse cTnnT protein distribution found in the lateral plate mesoderm (Figure 1D, arrows) ^13^. This event was followed by the establishment of the cardiac crescent-an epithelium-like structure reminiscent of a mesenchymal-epithelial transition and preceding the formation of the heart tube ^14, 6, 13^ (Figure 1D). We conclude that the *cTnnT-2a-eGFP* reporter faithfully identifies cardiomyocytes among a population of lateral plate mesodermal cell derivatives.

### Lineage analysis using a tamoxifen-inducible T/Bra reporter

Previous lineage tracing analysis revealed a temporal order in which different cardiac lineages arise within the mesoderm ^7 4^. To further assess the contribution of the temporal distinct mesodermal populations to the cardiac crescent and heart tube cell populations, we performed similar genetic tracing using an inducible *T^nGFP-CreERT^*^2^*^/+^* mouse, expressing *CreERT2* and nuclear localised GFP (nGFP) downstream of the endogenous *T* ^15^ in combination with the *R26R^tdTomato/+^* reporter and our novel cardiomyocyte *cTnnT-2a-eGPF* reporter (*T^nEGP-CreERT^*^2^*^/+^; R26R^tdTomato/+^*; *cTnnT-2a-eGFP*).

At ∼E7.75, the mesoderm is partitioned into three progeny compartments within the cardiac crescent: prospective endocardium, prospective myocardium, and prospective pericardium ^16, 6^. At later somite stages (E8.25), the cardiac crescent has generated the heart tube. The heart tube has an inverted Y shape, and the two arms of the Y -or inflows-, positioned inferiorly, are fated to become the atria, with the stem of the Y becoming the left ventricle (LV) and atrioventricular canal (AVC) ^17^.

Doses of tamoxifen (0.02 mg/bw) were administered at successive gastrulation stages ^7^. Early administration at E6+7h resulted in sparse tdTomato+ labelling of myocytes, endocardial, and pericardial cells in the cardiac crescent (n=3/3 embryos, Figure 1E-H), as well as in the endoderm. Later administration at E6+21h showed similar sparse labelling in most cases (n=7/10 embryos) (Figure 1Fii), while in some cases (n=3/10 embryos), no contribution to GFP+ myocytes cells was observed (Figure 1Fiii). As previously noted, variability in embryonic stages within litters at E6+21h likely explains these differences, with some embryos at more advanced stages having already downregulated *T/Bra* expression in cardiac crescent progenitors ^7^.

For both E6+7h and E6+21h tamoxifen regimens, a higher number of tdTomato+ *cTnnT-2a-eGFP+* myocytes were labelled compared to endocardial and pericardial progenitors (Figure 1H). This disparity suggests that the initial T/Bra-expressing mesoderm contains a greater number of myocyte progenitors. This finding likely explains the higher number of myocytes in the cardiac crescent relative to endocardial and pericardial cells. In some cases (n=5/13 embryos), we observed a large number of tdTomato-labeled GFP+ myocytes (between 57 and 127 cells) with rare or no corresponding tdTomato-labeled endocardial cells (between 0 and 8 cells) (Figure 1Giii). This observation supports the existence of independent progenitors that specifically contribute to myocytes in the cardiac crescent, but not to the endocardium ^2, 4, 18 (preprint)^. These results are consistent with a previous Foxa2 lineage tracing study, which identified independent *Foxa2+* progenitors contributing to ventricular myocytes, pericardium, and epicardium, but not the endocardium with a similar tamoxifen regimen ^7^.

When tamoxifen was administered at E7+7h, we observed a rare contribution of tdTomato+ cells to myocytes in the cardiac crescent (n=3/8 embryos, mean 10 tdTomato+ myocytes per embryo) and no contribution to endocardial or pericardial progenitors for all embryos analyzed (Figure 2Giii). At the heart tube stage, however, there was a strong contribution of tdTomato+ cells to inflow myocytes (>100 cells in all embryos analyzed, n=4/4 embryos), and inflow endocardial cells (39 cells on average, n=4/4 embryos) which are destined to form the atria during later development (Figure 1Ii-ii). This finding suggests an independent source of T/Bra-positive late mesodermal cells contributing to inflow myocytes and endocardium, as indicated in previous lineage tracing studies ^7,19^. Determining the clonal relationship between these cell populations will require tracing individual progenitors from their mesodermal origins to their final positions in the heart tube. As previously noted in a similar experiment ^7^, we also observed cases with sparse tdTomato+ endocardial cells in the ventricular regions, without any tdTomato+ cells in the ventricular myocyte GFP+ layer (n=2/4 embryos) (Figure 1Iii). This observation further supports the idea that independent sources exist for myocyte and endocardial progenitors.

**Figure 2.**
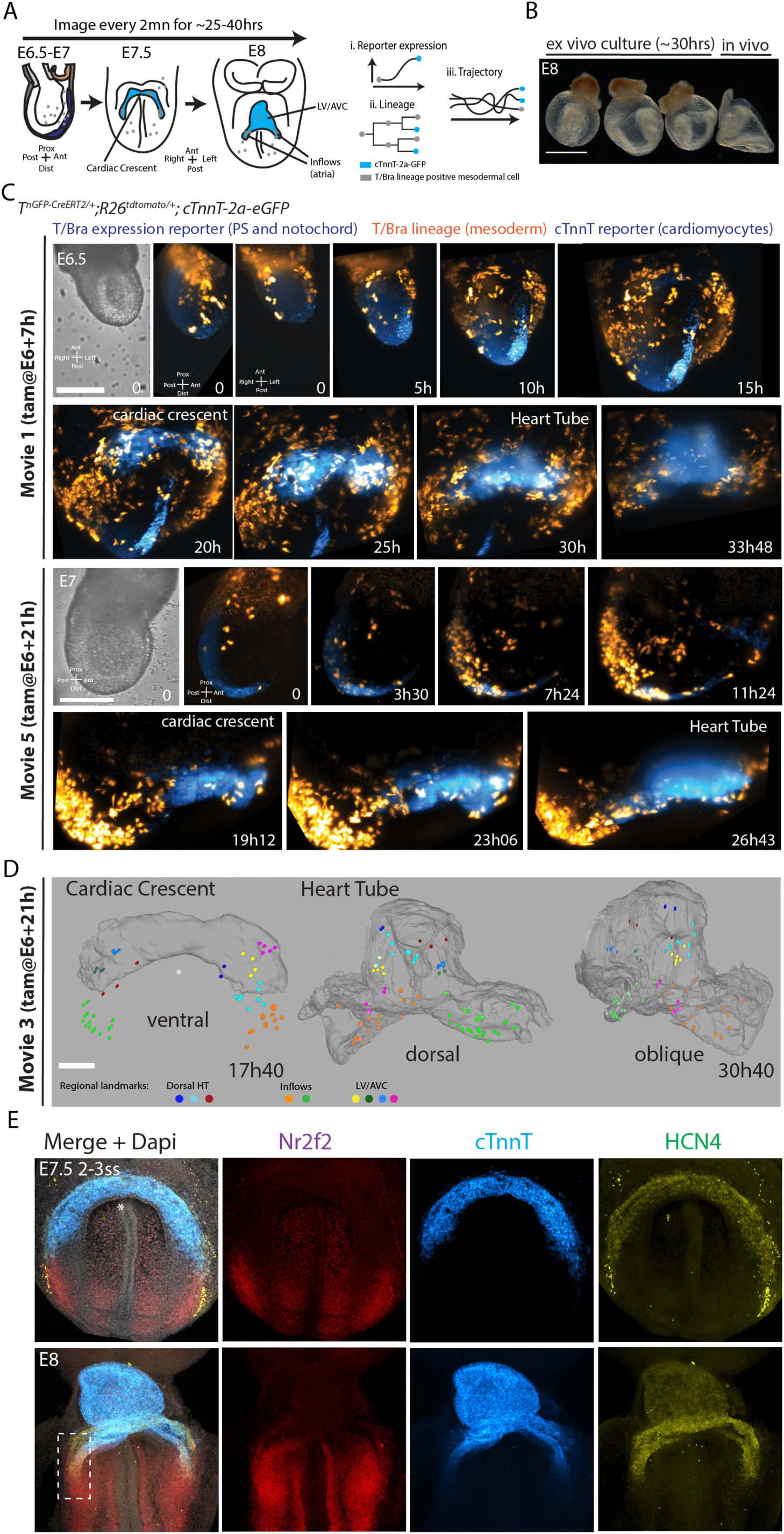
Live Analysis of Cardiogenesis from Gastrulation to Heart Tube Formation. (**A**) Schematic of cell tracking showing reporter expression, lineage relationships, and cell trajectories. (**B**) Example of embryos cultured ex vivo for 30h. (**C**) Time-lapse sequences of *T^nGPF-CreERT^*^2^*^/+^;R26R^tdTomato/+^; cTnnT-2a-eGFP* embryos after tamoxifen administration (0.02 mg/g) at the indicated times. Scale bar: 100 μm. (**D**) Fate mapping of the cardiac crescent. Cells are color-coded by their final position in the heart tube. Scale bar: 100 μm. (**E**) HCR in situ hybridization in E7.5-E8 embryos showing expression of HCN4, Nr2f2, and cTnnT, with the inflow region highlighted at E8.

### Establishing long term light-sheet live microscopy for cardiac lineage analysis

We employed live imaging and single-cell tracking, using the cardiomyocyte-specific cTnnT-2a-eGFP reporter, to reconstruct the lineage trees of mesodermal cells. Our objective was to identify the initial mesodermal progenitors contributing to the distinct cell populations within the heart tube of the gastrulating mouse embryo (Figure 2A). This approach required the culture and imaging of early-stage mouse embryos, spanning from the onset of gastrulation to the heart tube formation stage, a period of approximately 25 to 40 hours.

We found the Viventis LS1 open-top light-sheet microscope allowed the culture of early mouse embryos over long periods of embryonic development (>24 hours) ^20^. Incubation media was stable and could be exchanged during acquisition. A large media volume (∼1ml) improved embryonic viability for long-term imaging. Embryos cultured from E6.5 and for up to 40 hours developed normally; a cardiac crescent formed and generated a heart tube corresponding to E8 embryos (Figure 2B and Figure 2-Supplementary Figure 1 A).

To permanently label mesodermal cells and their progeny at a density suitable for live cell tracking, we used an inducible *T^2a-cre/ERT^*^2^ mouse combined with *R26^tdomato^* reporter and administered intermediate doses of tamoxifen (0.02mg/bw) at E5, E6+7h and E6+21h ^7^. We first administered tamoxifen earlier – (at E5) - and cultured embryos in tamoxifen-free culture media, from before the start of the gastrulation period and onset of T/Bra expression in the primitive streak and mesoderm – (at E6) -. In these conditions, no tdTomato expressing cells could be identified in the intra-embryonic mesoderm over ∼11 hours of live-imaging acquisition (Figure 2-Supplementary Figure 1B). This finding confirms that creERT2 activity in *T^2a-cre/ERT^*^2^ embryos requires *T/Bra* expression ^7^. In the absence of tamoxifen, rare tdTomato-positive cells were identified in only one embryo (not shown), confirming that tdTomato widespread mesodermal expression in *T^2a-cre/ERT^*^2^; *R26^tdTomato^* embryos requires tamoxifen.

We generated five light-sheet live-imaging datasets spanning 23 to 41 hours of mouse embryonic development from gastrulation – (E6.5-E7) - to heart tube stage (Figure 2C. and Figure 2 Supplementary Figure 2A and Videos 1-2). Embryos were imaged at 2 minutes intervals. Raw data amounts to 5-7 terabytes per experiment, representing up to half a million images. To correct for drift during acquisition, BigStitcher ^21^ was used to register the datasets in 4D as previously described ^6^.

Each movie contains up to ∼1200 time points, and a small percentage (<1%) of linkage inaccuracy between cells could lead to lineage misinterpretation that propagates over the course of the movie. Although automated cell tracking methods have seen major advances ^22,23,24^, it is still the case that achieving the level of precision necessary to reconstitute cell lineages over protracted periods remains challenging. To obtain accurate cell lineages, we manually tracked single *T/Bra* lineage positive cells by visualising them at successive time points from the beginning to the end of the movie using Massive Muti-view Tracker (MaMut) ^5^. We interrupted a track when it was impossible to identify the same cell across two successive time points unequivocally. A total of 227 mother cells were tracked for up to 5 generations resulting in 1299 descendants (Figure 3A).

**Figure 3.**
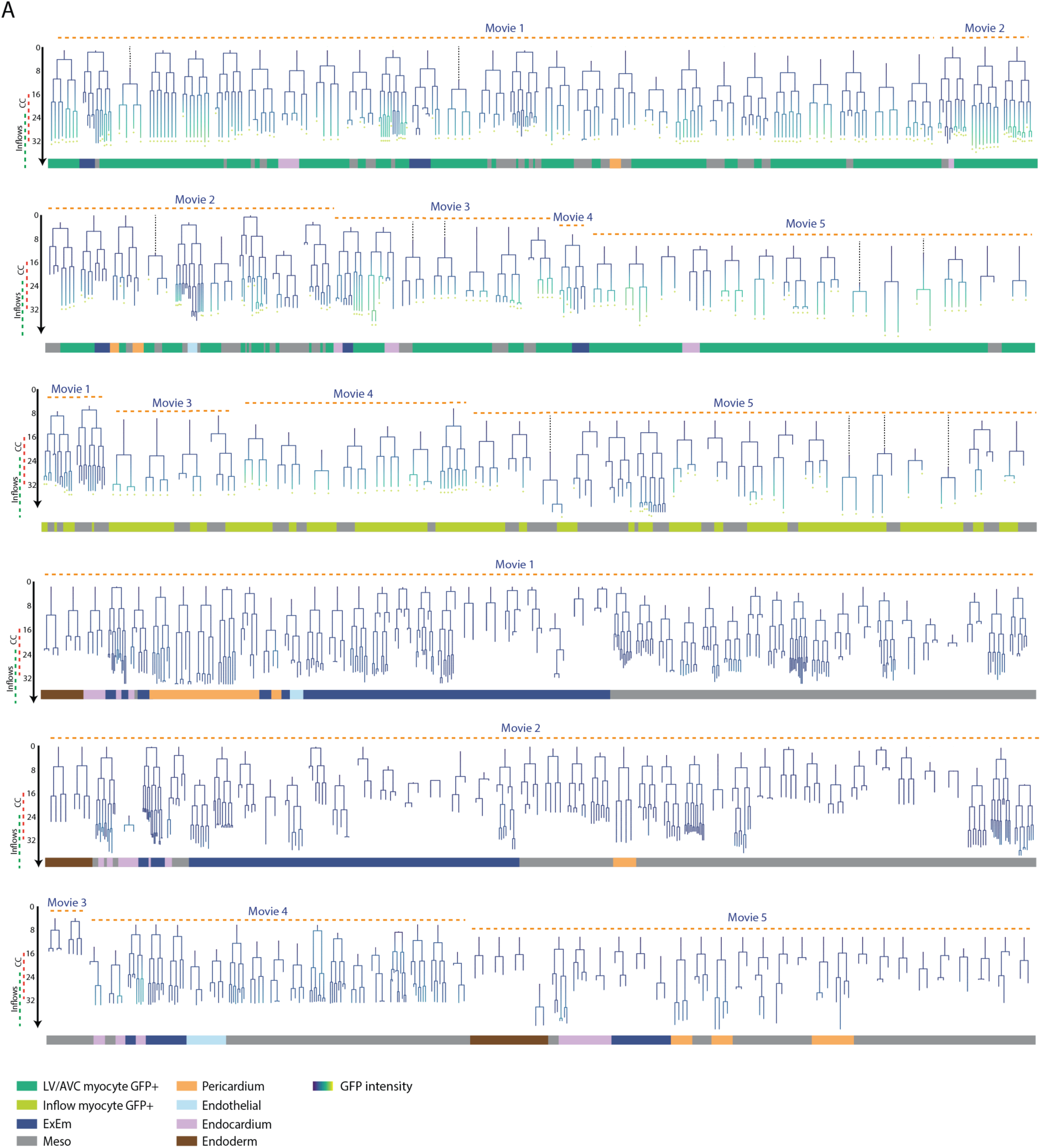
Reconstruction of the early mesodermal lineage tree. **(A**) Lineage trees of tracked progenitors. Cell types are shown at the endpoints, with GFP-positive descendants indicated by a yellow plus sign. Lineages are color-coded based on normalized GFP intensities. Grey, dashed lines indicate mother cells that could not be tracked back to their birth. Progenitors not contributing to any LV/AVC myocytes are shown in the bottom panel.

We determined the identity of the final daughters based on their location in the heart tube: within an inner endocardial layer ensuring the presence of a circulatory system, a myocardial layer formed by cTnnT-2a-eGFP+ cardiomyocytes and derived from the splanchnic mesoderm, and an outer layer derived from the somatic mesoderm called the pericardium (Videos 3-5). We could discriminate these cell types in our live-imaging datasets. Moreover, cardiomyocytes were further distinguished by their higher levels of *cTnnT-2a-eGFP* reporter expression (Figure 2-supplementary Figure 3A-F and Material and Methods). A substantial number of mother cells (n=111) produced at least one progeny whose fate could not be determined. In these cases, we could not determine if the mother cells were unipotent or multi-potent. This limitation may be due to some cells not reaching the GFP threshold required for classification as myocytes, likely influenced by their position in deeper embryo sections. Finally, the locations of the myocyte *cTnnT-2a-eGFP*+ cells within the LV/AVC and inflow regions of the heart tube indicated their fates. In what follows, we describe the lineage trees and timing for mesodermal progenitors’ specification into distinct cardiac lineages (Figure 3A).

### Heart tube morphogenesis

Initial analysis during the stages that the heart tube develops reveals a progressive propagation of myocyte differentiation and tissue folding along the anterior-posterior axis (Figure 2D). Myocyte differentiation is first triggered in the anterior regions that generate the cardiac crescent, and as tissue folding proceeds following epithelialization, the most anterior cells reposition to assume a more ventral position within the LV/AVC components of the heart tube. As differentiation and folding continue posteriorly, posterior cells contribute to the dorsal closure and inflow regions of the heart tube (see colour coded regional landmarks in Figure 2D and Video 6). Thus, heart tube morphogenesis occurs through a coordinated process of differentiation, epithelialization, and tissue folding, with the cardiac crescent initially forming the ventral aspect of the heart tube, while the inflow and dorsal aspects are progressively recruited to complete its formation.

I*n situ* HCR analysis further showed that the cardiac crescent and posterior mesoderm, destined to form the LV/AVC and inflow components of the heart tube respectively, exhibit distinct gene expression profiles (Figure 2E). HCN4, a first heart field marker ^25,26^, is expressed in both the prospective LV/AVC and inflow components of the heart tube, while Nr2f2, a marker required for atrial lineage specification ^27^, has its expression restricted to the posterior mesoderm and inflows myocardium in the heart tube (E7.5 and E8.5 in Figure 2E).

### Distinct mesoderm contributes to the heart tube and inflow myocardium

We next addressed the timing of LV/AVC and inflow myocyte lineage segregation. From our five datasets, we identified 91 progenitors contributing to at least one cTnnT-2a-eGFP+ myocyte (Figure 3A). Of these 91 progenitors, 61 contributed to the cardiac crescent, giving rise to 272 descendants, with 47 of these 61 progenitors traceable all the way to the heart tube. Additionally, 30 progenitors contributed to atrial cells, generating 84 descendants.

Previous clonal labelling of single mesodermal progenitors demonstrated the existence of precursors restricted to specific anatomical locations within the heart ^1,3,4,28^, while clonal induction at earlier embryonic stages typically generated larger clones that span multiple heart compartments, contributing to both heart fields ^3,4,28^. Consistent with these findings, our cell tracing at the mesoderm stage showed that most traced cells exhibit limited dispersion within the heart tube. Analyzing the progeny locations within the heart tube, we identified a clonal boundary at the junction between the LV/AVC and inflow myocyte compartments, suggesting that atrial and LV/AVC progenitors have distinct mesodermal origins (Figure 4Ai-iii and Figure 4-Supplementary Figure 1A-D). None of the clones contributed to both the LV/AVC and inflow myocyte compartments.

**Figure 4.**
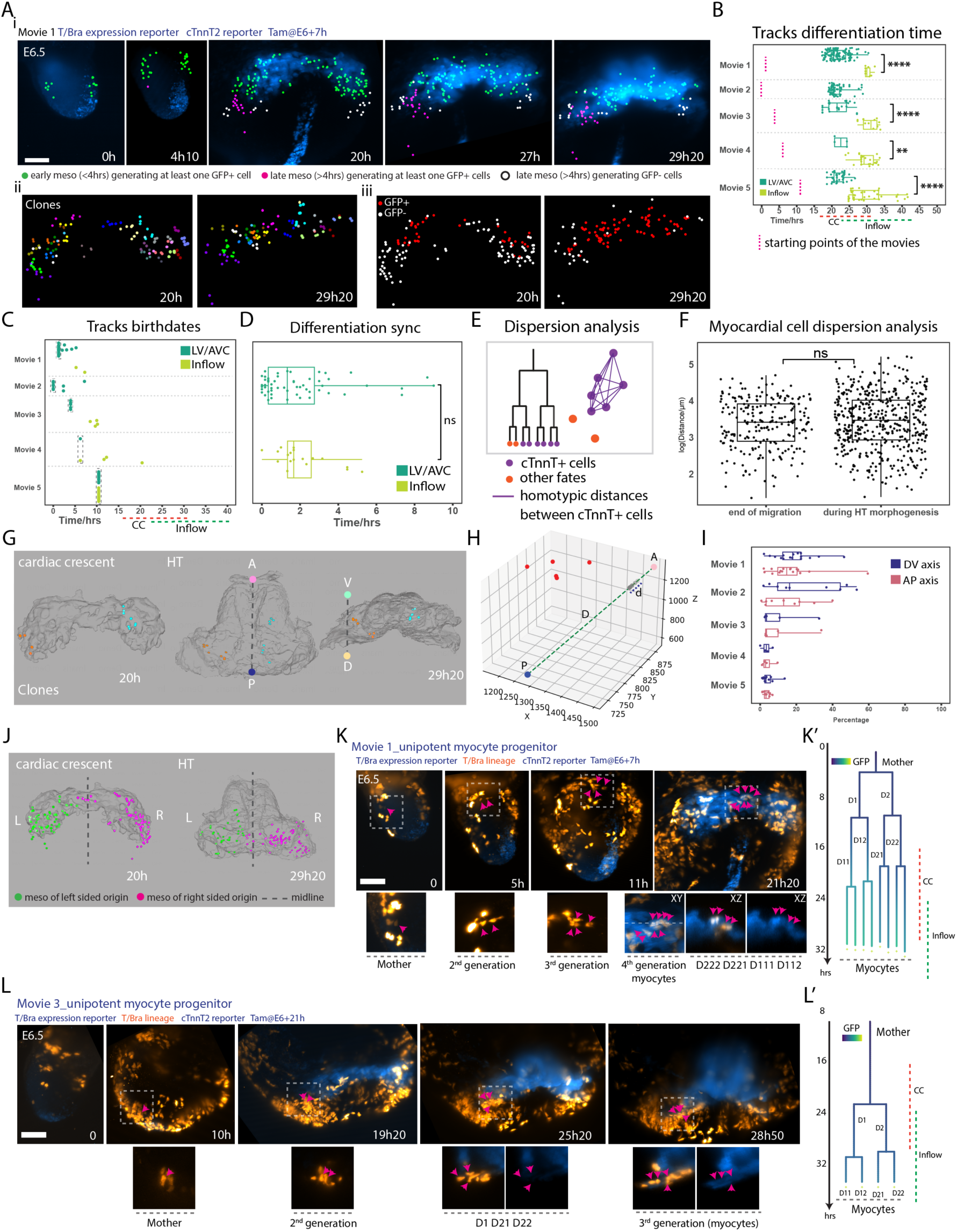
Independent LV/AVC and Inflow progenitors contribute to distinct regions of the heart tube. (**A**) (i) Fate maps showing early (green) and late (magenta) mesoderm contributing to cTnnT-2a-GFP+ progenitors; non-contributing late progenitors are in white. (ii) Each color represents a distinct clone. (iii) cTnnT-2a-GFP+ cells are in red, cTnnT-2a-GFP– cells in white. (**B**) Time points at which LV/AVC and inflow progenitors exceed the GFP intensity threshold. (**C**) Birth dates of progenitors contributing to cTnnT-2a-GFP+ LV/AVC and inflow myocytes. (**D**) Time between the first and last progenitor becoming cTnnT-2a-GFP+ in myocyte lineages. (**E**) Schematic of dispersion analysis in lineages. (**F**) Comparison of cell-cell dispersion for cTnnT-2a-GFP+ myocytes at migration’s end (defined by the mean LV/AVC differentiation time in each movie) and during heart tube formation (41.4 µm ± 31.9 (SD), n=293 at the end of the migration period; 45.1 µm ± 34.7 (SD), n=597, after heart tube formation (p=0.22). (**G-H**) Example of myocardial clones dispersed along anteroposterior (AP) and dorsoventral (DV) axes. (**I**) Dispersion along AP and DV axes across movies. Boxplots show median, 25th/75th percentiles, and whiskers for minimum/maximum values. (**J**) Locations of mesodermal progenitors from left/right nascent mesoderm contributing to cTnnT-2a-GFP+ cells. (**K-L**) Time-lapse images and lineage trees of LV/AVC (K) and inflow (L) progenitors, colored by normalized GFP intensity. Scale bar: 100 μm. Statistical analyses: Mann-Whitney U test.

In most cases, clones were confined to specific dorso-ventral (DV) and antero-posterior (AP) positions, with 35 out of 44 clones spreading less than 20% along the longest AP axis of the LV/AVC heart tube region, and 32 out of 44 clones spreading less than 20% along the longest DV axis (Figure 4H-I and Video 7). In rare instances, we observed clones spreading across both the ventral and dorsal regions, as well as the anterior and posterior regions, particularly when tracked from earlier stages of gastrulation. These clones dispersed across 40% to 60% of the entire AP (n=1/44) and DV (n=4/44) axis lengths (Figure 4G and I; Movie 1 and 2). In even rarer cases (n=1/44), we observed cells crossing the midline, with a clone spanning both the left and right sides of the cardiac crescent and heart tube (Figure 4J and K, indicated by red arrows).

Analysis of distances between sister cells shows that, on average, clones do not spread significantly further during the stages of heart tube formation (Figure 4E-F). This finding indicates that most clonal dispersion occurs during earlier cell migration, whereas the heart tube formation is primarily driven by tissue-scale processes like epithelial deformation and folding, rather than individual cell movements or intercalation ^6,13,14^. Consequently, clonal spreading is minimal during heart tube formation.

LV/AVC progenitors are born first and differentiate into *cTnnT-2a-eGFP*+ myocytes before other cardiomyocytes. This establishes the initial cardiac crescent within a ∼ 15-hour period with the first LV/AVC progeny differentiating at 16 hours and the last at 31 hours. Atrial progenitors are born later, differentiate later -from 24 hours to 42 hours -, and are recruited to posterior regions during the folding of the cardiac crescent into the heart tube. This event establishes the inflows (Figure 4 A-C and K-L).

Myocytes developed concurrently within each lineage (Figure 4D). The time intervals between the first and the last daughter to transition into cTnnT-2a-eGFP+ myocytes were similar on average between the LV/AVC and atrial lineages (means of 2.0 hours and 2.1 hours for LV/AVC and atrial lineages respectively). Notably, the differentiation timing varied among lineages, with some displaying greater synchrony than others. For instance, in 5 out of 55 lineages, the mother cell generated LV/AVC cTnnT-2a-eGFP+ myocyte daughters in more than 5 hours. In contrast, in 25 out of 55 lineages, all daughters transitioned into cTnnT-2a-eGFP+ myocytes in less than 1 hour.

Together, the live-imaging analysis shows that the heart tube is established by at least two sets of independent LV/AVC and inflow myocyte progenitors generated from early and late mesoderm respectively and differentiating into myocytes at different embryonic stages. This observation is in line with the previous hypothesis that atrial and LV/AVC compartments have distinct spatial and temporal origins during gastrulation in the mouse ^7–9^.

### Identification of unipotent, bipotent and tripotent mesodermal progenitors

We next investigated whether the mesoderm is predominantly composed of multipotent cells, which have the potential to differentiate into various cell types, or a heterogeneous mixture of cells that are already committed to specific fates ^29^.

Among the five embryos analyzed, we found that the majority of progenitors tracked within the early proximal mesoderm contributed to the LV/AVC (n=61), whereas endocardial (n=18) and pericardial (n=13) progenitors were less common in the mesoderm. This disparity, in contribution to the three cell types, -also shown by lineage tracing experiments in a larger cohort of embryos (Figure 1H)-likely explains the higher number of myocytes compared to endocardial and pericardial cells within the cardiac crescent, indicating that a larger number of LV/AVC progenitors is necessary to account for these differences.

A substantial number of mother cells produced at least one progeny whose fate could not be determined (n=111) (see Figure 5C for examples). In these cases, we were unable to determine whether the mother cells were unipotent or multipotent. Focusing on progenitors for which we could determine potency, our analysis revealed a predominance of unipotent progenitors (n=98, generating 728 descendants) compared to bipotent/tripotent progenitors (n=18, generating 302 descendants). We identified 29 unipotent mother cells generating only LV/AVC myocytes and 12 bipotent/tripotent progenitors contributing to at least one LV/AVC myocyte descendant. Thus, approximately a third (12/41) of the mother cells contributing to LV/AVC myocytes, for which we could determine potency, were bipotent or tripotent.

**Figure 5.**
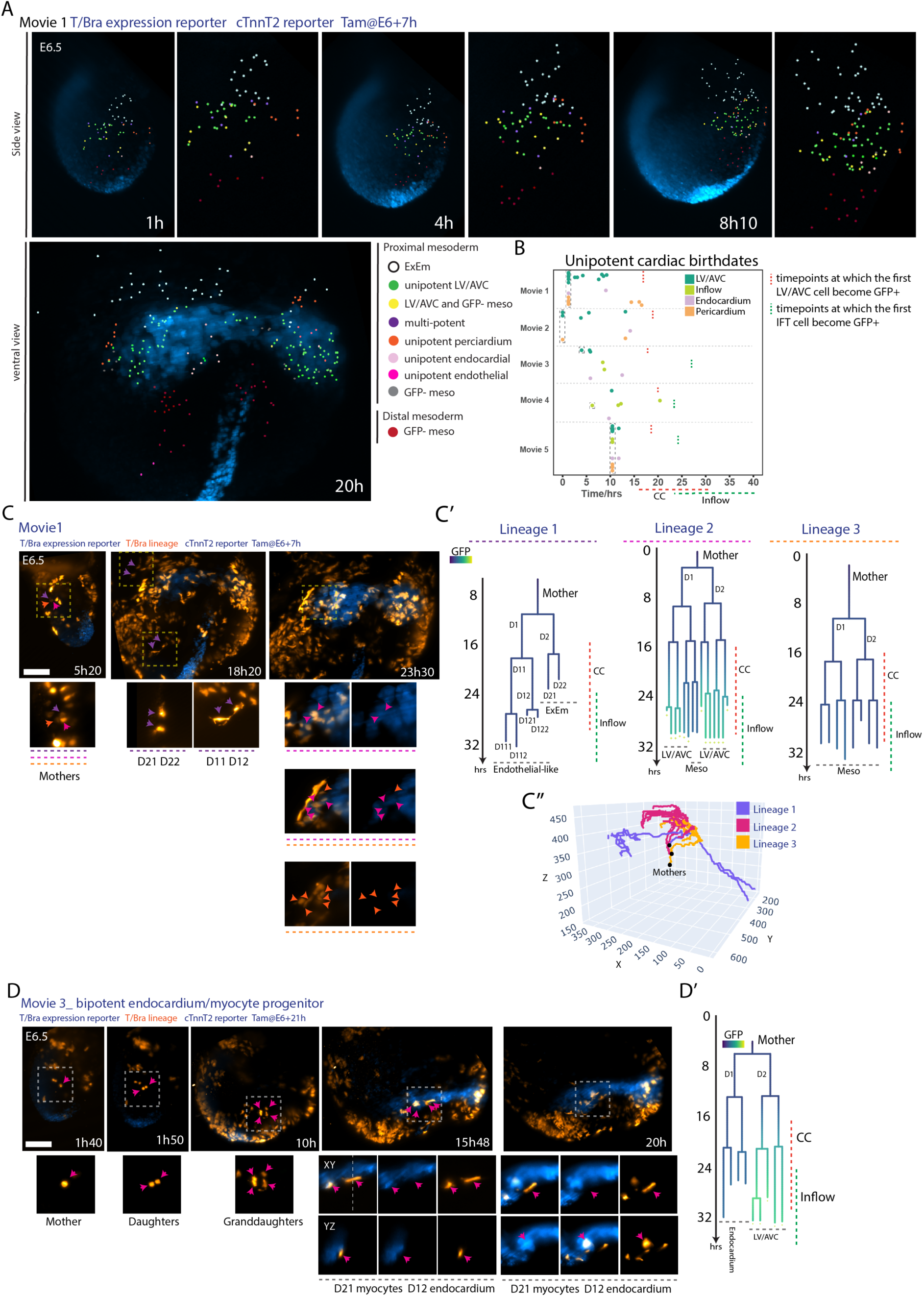
Fate Mapping of Cardiac Progenitors at the Gastrula Stage. (A) Cells are color-coded by fate. (**B**) Birth dates of unipotent progenitors for LV/AVC, inflow myocytes, endocardium, and pericardium. Cells present at the start of the movie are highlighted in a grey-dotted box. (**C**) Time-lapse sequences of *T^nGPF-CreERT^*^2^*^/+^;R26R^tdTomato/+^; cTnnT-2a-eGFP* embryos showing three nearby progenitors contributing to extraembryonic mesoderm (ExEm), endothelial-like cells, LV/AVC, and undefined mesodermal fates. Corresponding lineage trees are color-coded by normalized GFP intensity (C’), with arrows indicating tracked cells. (C’’) 3D trajectories of the three lineages. Scale bar: 100 μm. (**D**) Time-lapse of *T^nGPF-CreERT^*^2^*^/+^;R26R^tdTomato/+^; cTnnT-2a-eGFP* embryos showing a bipotent progenitor. Corresponding lineage trees are color-coded by normalized GFP intensity (D’), with arrows indicating tracked cells. Scale bar: 100 μm.

We could not determine whether certain types of bipotent progenitors were more prevalent than others. Instead, we observed a diversity of bipotent progenitor cell types, including LV/AVC-endocardial (n=4), LV/AVC-ExEm (n=4), LV/AVC-pericardial (n=2), and LV/AVC-endothelial-like (n=1), as well as endocardial-ExEm (n=3) and pericardial-ExEm (n=1) bipotent progenitors and one tripotent progenitor contributing to LV/AVC myocytes, endocardium, and extra-embryonic mesoderm (Figure 3A, Figure 5C-D, Figure 5 Supplementary Figure 1A-C, and Figure 6 Supplementary Figure 4F, Figure 7, Supplementary Figure 3A-B). One additional bipotent progenitor generated highly migratory endothelial-like progeny located in both the embryonic and extra-embryonic mesoderm (Figure 5C’).

**Figure 6.**
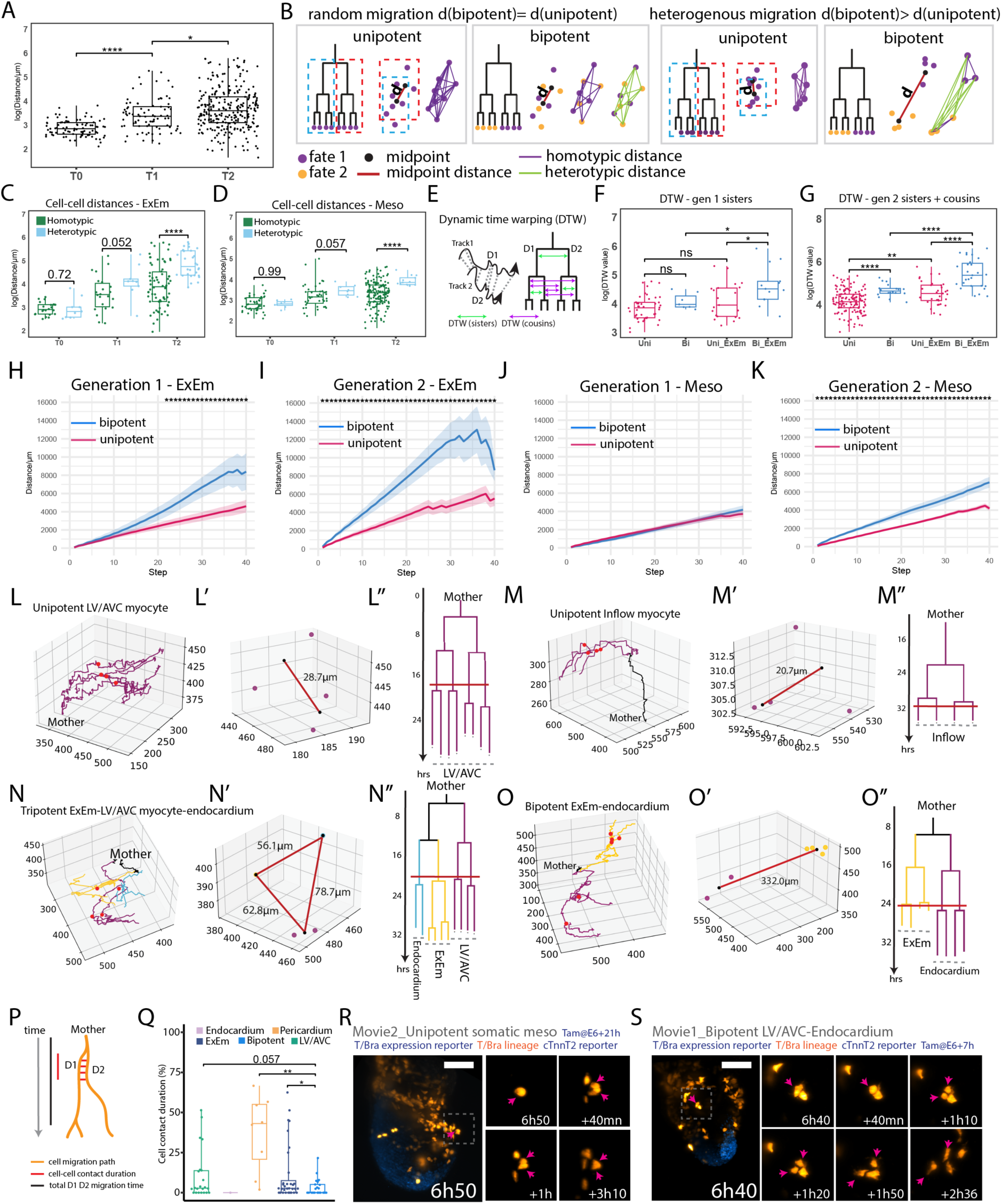
Cell migration analysis in lineages reveals hidden patterns. **(A)** Dispersion analysis of daughter and granddaughter cell distances in cardiac lineages at T0, T1, and T2. **(B)** Hypothesis schematic: random migration predicts equal dispersion for unipotent and bipotent progenitors; heterogeneous trajectories predict less dispersion for unipotent progenitors. “Midpoint distance” is the Euclidean distance between the midpoints of D1’s and D2’s daughters. **(C-D)** Dispersion analysis comparing homotypic and heterotypic distances in lineages with (C) and without (D) ExEm cells at T0, T1, and T2 (log scale). (**E**) Dynamic time warping (DTW) analysis of migration path similarities. **(F)** DTW scores comparing D1 and D2 sister tracks up to their next division for unipotent vs. bipotent fates (log scale). **(G)** DTW scores for 2nd generation descendant tracks (sisters [D11-D12, D21-D22] and cousins [D11-D21, D11-D22]) based on shared or distinct fates (log scale). **(H-K)** Mean DTW values over time for D1, D2 (generation 1) and D11, D12, D21, D22 (generation 2) daughters from unipotent and bipotent progenitors, with ExEm cell contribution (H-I) or without (J-K). Shaded areas indicate SE; significant differences for each steps are marked with a star. **(L-O)** Examples of trajectories and corresponding midpoint distances (L’-O’) and lineage trees (L’’-O’’). Red lines mark sampling points for midpoint analysis. **(P)** Cell-cell contact duration analysis: ratio of contact time to track duration within the first 16 hours. Short tracks (<4 hours) excluded. **(Q)** Proportion of time sister cells with similar or distinct fates remain in contact. **(R-S)** Time-lapse images of unipotent pericardial (R) and bipotent LVAVC/Endocardium (S) progenitors. Statistical analyses: Mann-Whitney U test.

**Figure 7.**
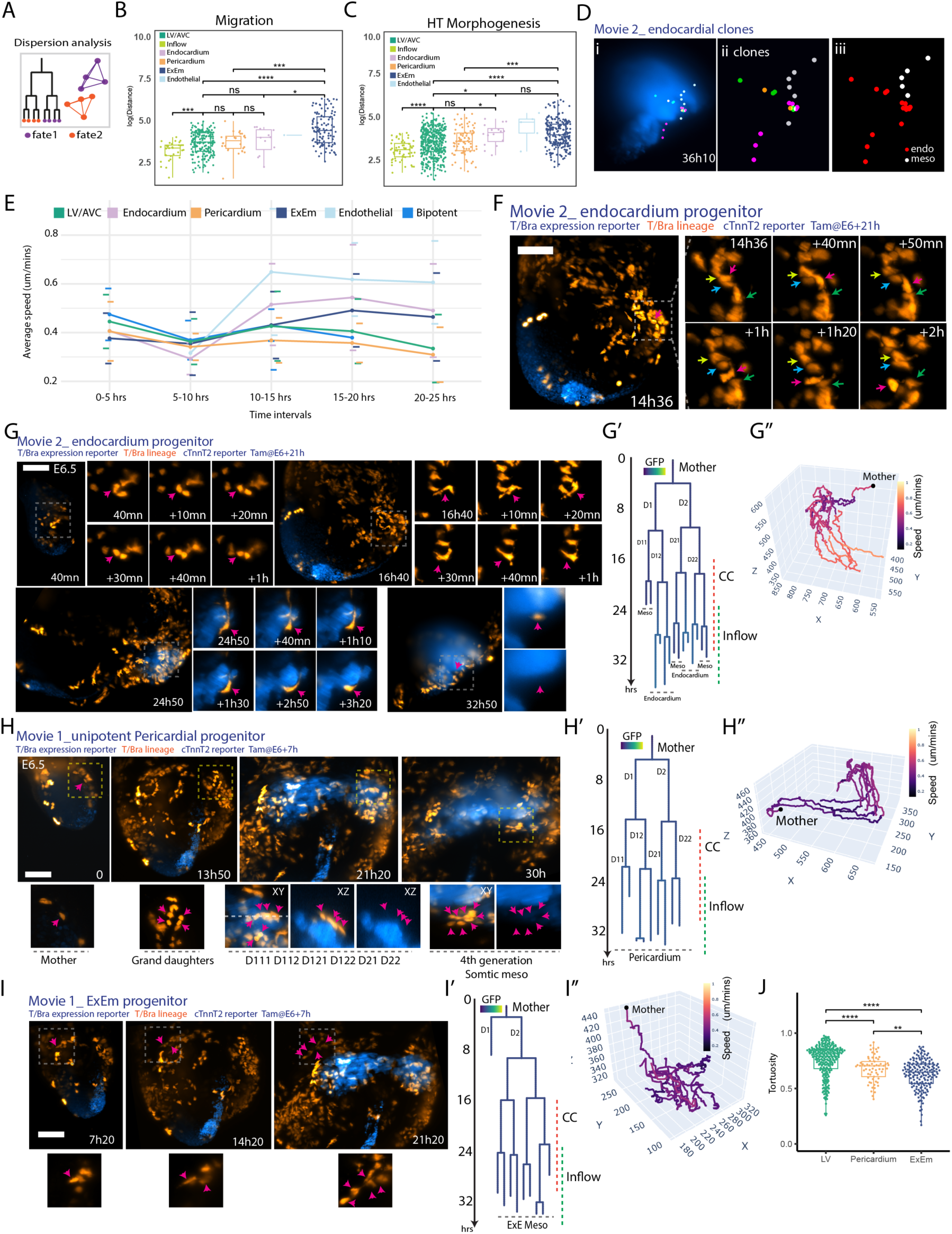
Analysis of Mesodermal Cell Behaviour. (**A**) Summary diagram of dispersion analysis for sister and cousin cell distances with similar fates. (**B-C**) Cell dispersion at migration end (B) and during heart tube formation (C). (**D**) (i-ii) Endocardial clones, each in a different color. (iii) Endocardial cells highlighted in red. (**E**) Mean cell speeds per fate calculated across 5-hour intervals (see Figure 6 Supplementary Figure 1A-G). (**F-I**) Time-lapse images of endocardial (F-G), unipotent pericardial (H), and ExEm (I) progenitors. Corresponding lineage trees, colored by GFP intensity, in G’, H’, I’, and 3D speed plots in G’’, H’’, I’’. Arrows in G-I indicate cells in lineage trees G’, H’, I’. (J) Tortuosity analysis comparing ExEm and pericardial cells during heart tube formation stages. LV/AVC: left ventricle/atrioventricular canal, HT: heart tube. Statistical test: Mann-Whitney U test.

**Figure 8.**
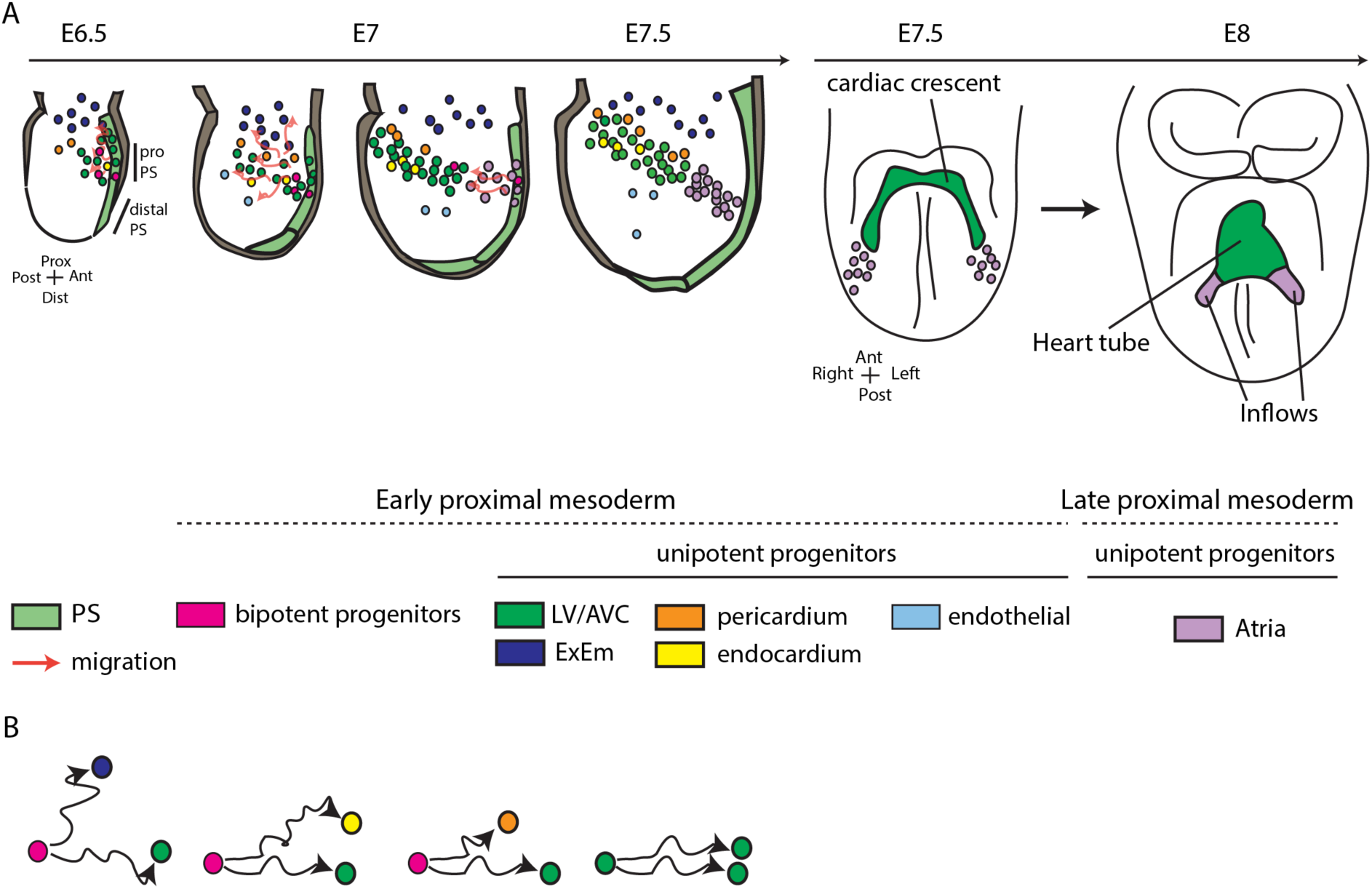
Working Model of early cardiac development. **(A)** Early proximal mesodermal cells are rapidly become committed to specific cardiac fates as they initiate migration towards specific embryonic regions. LV/AVC and Atrial progenitors are generated from early and late mesoderm, respectively. LV/AVC progenitors differentiate first into myocytes and establish the cardiac crescent. Atria progenitors differentiate later and generate the heart tube’s inflows. (**B**) Pair of sisters sharing the same fate have more identical migration paths than sisters with distinct fates. PS: primitive streak. LV/AVC: left ventricle/atrioventricular canal. ExEm: Extra-embryonic mesoderm.

Our lineage analysis led us to examine the initial locations of unipotent and multipotent progenitors within the early proximal mesoderm. We observed that LV/AVC, pericardial, and ExEm progenitors showed a tendency to localize in separate regions (Figure 5A). Notably, unipotent pericardial progenitors were initially situated in anterior regions, consistent with a recent analysis in zebrafish indicating that pericardial cells originate from locations distinct from the heart field ^30 (preprint)^. However, we found that initial cell positions did not strictly correlate with their eventual fates. For instance, three neighbouring cells could migrate to different regions of the embryo and adopt diverse fates (Figure 5C-C’’). These results suggest that precise positional information dictating cell fate at cellular resolution may not be present in the early proximal mesoderm. Moreover, bipotent and tripotent progenitors were intermingled with unipotent LV/AVC progenitors, lacking any discernible spatial pattern (Figure 5A).

Consistent with previous live-imaging analyses ^6,7^, the distal mesoderm—already present in the early stages of the gastrulating embryo (Figure 5A, dark red cells at 1h)—migrated to more medial locations known to give rise to progeny that contribute to the right ventricle, outflow tract, and branchiomeric muscles ^31–32^. No T/Bra lineage-positive medial mesoderm was identified in movie 5 (Figure 2C).

In the late mesoderm, we identified 13 unipotent atrial myocyte progenitors out of the 30 progenitors contributing to the cTnnT-2a-eGFP+ atrial myocytes. Longer tracks encompassing later stages will be necessary to determine whether the remaining progenitors contribute exclusively to cTnnT-2a-eGFP+ atrial myocytes or if they also give rise to additional lineages. Moreover, we found additional progenitors contributing exclusively to cTnnT-2a-eGFP-daughters, located in the inflow regions of the heart tube and the posterior lateral plate mesoderm (n=24 lineages, identified as meso GFP-in Figure 3A). Longer imaging periods will be required to determine the identity of these cells. A subset of these cells displayed spindle-like shapes and were identified as endothelial-like cells (n=4, Figure 3A).

Together, the live-imaging analysis of lineages reveals that early mesodermal cells harbour plasticity and diversity of fates during gastrulation ^1^. However, their ability to alternate fates appears to diminish rapidly. In all cardiac lineage trees that displayed two or three fates (n=18), progeny became lineage-restricted early, during migration, before the onset of cTnnT-2a-eGFP+ expression in the embryo (Figure 5B). Of the 18 bipotent progenitors, 12 became unipotent after the first generation, 5 after the second generation and 1 after the third generation (Figure 3A and Figure 5B). These findings are consistent with previous clonal analyses, suggesting that early mesodermal progenitors are rapidly specified into discrete fates shortly after the initiation of gastrulation ^3, 4, 1^.

### Migration analysis in lineages reveals hidden patterns of cell migration

Previous live analysis of cell trajectories during gastrulation revealed apparently chaotic individual cell movements during migration ^6^. Consistent with this analysis, we found that mesodermal cells dispersed extensively during migration (Figure 6A). We analyzed distances between the first two daughters (coordinates were taken 10 minutes after the first cell division - Timepoint 0 -, and last time point before the daughters’ subsequent cell division - Timepoint 1 -) and granddaughters (final time point at which all granddaughter cells exist; we only considered branches lasting at least 4 hours into the cell cycle to allow sufficient cell migration - Timepoint 2 -) in each lineage. Distances between daughters and granddaughters gradually increased, reaching considerable distances within a single lineage (up to 331 μm) (Figure 6A). We noted, however, that distances were highly heterogenous; a proportion of the progeny generated less dispersive daughters at T2, with separating distances of less than 25 μm between them (80 out of 268 calculated distances across 44/59 lineages). One possibility for the observed heterogeneity in these distances is that the daughters generated by unipotent progenitors exhibit less dispersive migratory paths than those generated by bipotent progenitors (Figure 6B). To test this hypothesis, we analyzed cell movements in lineages, taking advantage of our lineage analysis from the live-imaging data.

Distances between sister cells can be large in non-cardiac lineages, sometimes exceeding 300 μm (n=3). Daughters and granddaughters from bipotent progenitors generating both ExEm and cardiac fates had greater distances than those from unipotent ExEm progenitors (Figure 6C at T2). Similarly, in the cardiac crescent, bipotent progenitors contributing to myocyte, endocardial, or pericardial fates (but not to the ExEm) produced daughters with greater distances than unipotent progenitors (Figure 6D at T2). While no differences were seen immediately after division (Figure 6C, D at T0), as development progressed, sisters with the same fate remained more closely positioned than those with distinct fates (Figure 6C, D at T2). This finding suggests unipotent progenitors produce more closely positioned cell descendants than bipotent ones.

These differences might be due to how similar or different cells’ migration paths are. To explore this possibility, we analyzed whether sisters sharing the same fate migrated closer together compared to those with divergent fates, using dynamic time warping (DTW) to account for temporal shifts in cell behaviour (Figure 6E, Materials and Methods). We calculated the cumulative DTW distances, a distance measure that assigns small distances to similar paths that are temporally unaligned (Figure 6E).

As expected, unipotent ExEm progenitors had lower DTW distances than bipotent progenitors, which produced cells contributing to both ExEm and one of the cardiac cell types (myocardium, endocardium, or pericardium). This divergence was evident from the first generation, indicating early divergence in the migratory paths of cells contributing to ExEm and cardiac fates (Figure 6F-G, N-O and Figure 6 Supplementary Figure 1H-I). For progenitors generating cardiac fates but not ExEm, unipotent progenitors also produced sisters with lower DTW distances compared to bipotent progenitors, becoming significant by the second generation suggesting a more gradual divergence in migratory paths (Figure 6F-G and Figure 6 Supplementary Figure 1F-G).

In 49 out of 168 cases, unipotent progenitors generated daughter cells with distinct migratory trajectories (log DTW > 4.5 at the second generation, including 32 non-ExEm progenitors). Conversely, in 5 out of 45 cases, bipotent progenitors produced sister cells that followed similar paths but adopted different fates (log DTW < 4.5 at the second generation, including one ExEm-contributing progenitor). These findings suggest that sister cells can diverge in migratory paths while sharing the same fate or follow similar paths but adopt different fates (Figure 6 Supplementary Figure 1A-B and J). However, in the majority of cases, sisters with the same fate exhibited notably similar migration patterns (Figure 6L-M and Figure 6 Supplementary Figure 1C-E).

A permutation test with 100,000 iterations confirmed that unipotent progenitors produced sisters with more similar trajectories than bipotent progenitors (p = 9.9e-07) (Figure 6 Supplementary Figure 2A-D and Material and Methods). Plotting DTW values over time showed that sisters with the same fate maintained similar migratory paths throughout, and any observed similarity is not attributed to systematic smaller DTW-contributions towards the end of the trajectory. By contrast, sister cells with distinct fates exhibited divergent migratory behaviours, with those contributing to ExEm showing a rapid and progressively accelerating divergence (increasing DTW values), while non-ExEm cardiac progenitors displayed a more gradual differences in paths (Figure 6H-K).

The observed similarity in migratory paths of sister cells with shared fates led us to investigate whether these cells retain contact longer post-division. We quantified this by measuring the ratio of contact duration to total tracking duration for generation 1 (Figure 6P, Figure 6 Supplementary Figure 3A). Results revealed that bipotent progenitors produced daughter cells with the shortest contact duration, displaying robust separating behaviours (Figure 6Q-S). By contrast, unipotent pericardial progenitors maintained the longest contact duration, suggesting sustained interaction, while LV/AVC unipotent progenitors showed a similar trend, albeit not statistically significant. These findings raise the possibility of correlation between progenitor fate and early contact behaviour.

### Emergence of distinct mesodermal cell behaviours during gastrulation

We next determine if distinct cell fates adopt different dispersion behaviours and when these differences emerge in the mesoderm. Initial dispersion analysis at the end of the migration period showed no significant differences between LV/AVC, pericardium, and endocardial fates. However, atrial sister cells clustered more closely, while ExEm cells were more dispersed (Figure 7A, B). During stages of heart tube formation, endocardial cells displayed greater dispersion than LV/AVC and pericardial cells, resembling the behavior of ExEm cells (Figure 7C). Consistent with this observation, endocardial clones can spread in both the ventricular and inflow component of the heart tube (Figure 7D). This finding suggests that endocardial cells adopt distinct migratory behaviours at later stages, leading to their further dispersion during heart tube formation. Supporting this observation, a concurrent analysis revealed behavioral divergence in endothelial/endocardial precursors, with endothelial progenitors exhibiting higher speeds^18 (preprint)^.

These findings led us to investigate cell speed patterns according to cell fate using our annotations. After aligning the movies to a unified timeline (Figure 4B), we analyzed cell speed in 5-hour intervals, revealing distinct trends. During the 0-5 hour interval, ExEm progenitors exhibited the slowest speeds ^6, 10^, while bipotent, pericardial, and LV/AVC progenitors showed similar speed. Endocardial cells initially migrated slowly but showed a marked increase in speed during the 10–15 hour interval, whereas pericardial progenitors exhibited a decrease in speed over the same period (Figure 7E and Figure 7-suplementary Figure 1A-G). At these stages, endocardial cells began transmigrating through the mesoderm, preceding cTnnT-2a-eGFP+ reporter activation, and adopting a spindle-like morphology (Figure 7F). Some were already positioned near the endoderm before cTnnT-2a-eGFP+ activation in both the LV and atrial regions (Figure 7 Supplementary Figure 2A, D, n=22). From around 25 hours, other endocardial cells crossed the cTnnT-2a-eGFP+ myocardium, contributing to the left ventricle (LV) and atrial components of the heart tube (Figure 7 Supplementary Figure 2B and C, Figure 7G, n=13). Similarly, endothelial-like cells exhibited spindle-like shapes and increased their speeds during the 10-15 hour interval, aligning with the timing of endocardial cell acceleration and cell shape changes (Figure 7E, Figure 7 Supplementary Figure 3A and Figure 7 Supplementary Figure 1A and G), consistent with those reported in Sendra et al. 2024 ^18 (preprint)^.

Finally, ExEm progenitors showed a significant increase in speed and higher tortuosity during the 15-20 hour interval, indicating more meandering and chaotic migratory paths (Figure 7I and Figure 7 Supplementary Figure 3B). In contrast, LV/AVC and pericardial cells displayed more directed movements, aligning with the onset of epithelialization and tissue folding in the cardiac crescent to form the heart tube, with pericardial tissue largely following these directed movements taking place at these stages ^6^.

## Discussion

Our findings illustrate the migration of cardiac mesodermal lineages during gastrulation. Using live-imaging and single cell tracking, we reconstructed cardiac mesodermal lineages and the migratory paths of cells over extended periods encompassing gastrulation and heart tube morphogenesis (∼40 hours). Culturing embryos in larger volumes of media culture using an open top light sheet microscope and in-house preparation of high-quality rat serum ^33, 34^ were critical for these experiments.

Our live-imaging suggests that the cardiac crescent (or first heart field -FHF- and juxta-cardiac field partially overlapping the FHF) ^25,26,35,36^ is predominantly destined to contribute to the LV/AVC. In contrast, the atria arise at different times during gastrulation ^1,4,7–9^. These findings suggest that an early segregation of the ventricular and atrial cells has been conserved during evolution; an early segregation of these progenitor populations was previously shown at single cell resolution in the zebrafish ^37–40^. They also align with *in vitro* differentiation experiments demonstrating that modulating pathways known to induce mesoderm can generate molecularly distinct mesoderm favouring the generation of ventricular or atrial-like cardiomyocytes respectively ^41–45^.

The limited initial spread of T/Bra lineage positive clones suggests that mesodermal cells quickly become confined to specific locations during gastrulation ^4, 3, 28,^ while large clones spanning multiple heart compartments may result from earlier induction events, possibly at the epiblast stage ^28^. However, this restricted distribution does not necessarily imply early segmental regionalization within a strictly predetermined mesoderm. Additional clonal spreading ^46, 47^, may occur during later stages as the heart undergoes oriented growth, reshaping and looping. Our live-imaging did not cover these later stages, leaving this possibility unconfirmed.

Our analysis showed that most early proximal mesoderm progenitors contributed to the LV/AVC, while fewer gave rise to endocardial and pericardial cells. This imbalance likely explains the higher number of myocytes compared to endocardial and pericardial cells in the cardiac crescent, suggesting that not all progenitors in the early proximal mesoderm are equivalent. Instead, specific biases in cell fate may be regulated to produce the correct cell types in the right proportions.

Our analysis revealed that the early proximal mesoderm contains bipotent progenitors capable of generating LV/AVC, pericardium, endocardium, and extraembryonic mesoderm. However, many mother cells produced at least one progeny whose fate could not be determined, making it difficult to confirm their potency and accurately estimate the incidence of bipotent cells due to challenges in reconstructing complete lineage histories. Addressing these limitations will require future studies to use more advanced microscopy techniques with higher contrast-to-noise ratios for improved visualization in deeper tissue layers.

Despite these challenges, our reconstruction of 227 early mesodermal lineage trees across up to five generations from five live-imaging datasets of gastrulating mouse embryos provides valuable insights into cell potency. Among the progenitors with confirmed potency, we identified 12 bipotent/tripotent and 29 unipotent progenitors contributing to LV/AVC myocytes. These findings suggest that while many mesodermal progenitors are committed to a myocyte fate, a substantial proportion of bipotent progenitors also exist. These bipotent progenitors became rapidly restricted into unique cardiac fates during migration - prior to the establishment of the cardiac crescent and onset of myocyte differentiation. The observation of short-lived multipotent cardiac progenitors is consistent with clonal analysis results of *Hand1+* and *Mesp1+* mesodermal progenitors ^1,3,4^.

Previous migration analysis noted opposing cell density and motility gradients in the mesoderm ^6^. According to this model, cells continually exchange neighbours and disperse widely until their movements gradually diminish, eventually settling in positions and fates as gastrulation concludes. Our live-imaging analysis builds upon these findings, offering an extended evaluation of the migratory paths of cells in relation to their future fates in the heart tube. The analysis revealed that progenitors contributing to LV/AVC and atrial myocytes remain as separated cell populations throughout migration, establishing two distinct progenitor domains in the heart tube without mixing.

During development, the pericardial and myocardial layers, along with the underlying plexus of elongated endocardial cells, emerge in close proximity within the cardiac crescent ^6,16^. The migratory paths of progeny contributing to these three distinct cardiac fates appear more predictable than previously understood, with sister cells sharing the same fate often following parallel trajectories over extended periods. Bipotent progenitors, particularly those giving rise to ExEm, displayed the fastest and most widely dispersed migration patterns among their daughter cells. In contrast, bipotent progenitors contributing to endocardial, LV/AVC and pericardial cells showed a more gradual divergence in migratory paths. This divergence was characterized by a late-stage increase in speed for endocardial cells, while pericardial cells exhibited a concurrent decrease in speed relative to LV/AVC progenitors. These gradual trajectory shifts likely reflect late-stage cellular behavioural changes associated with the process of fate commitment during gastrulation.

Future research will need to explore the potential mechanisms restricting migratory paths and determine whether early mesodermal progenitors display similar migratory behaviours due to shared initial internal states or a cell-specific response to environmental cues. Previous studies in zebrafish indicated that G-protein-coupled receptor signalling, a hallmark of chemokine signalling, regulates heart progenitor movements during gastrulation ^48–50^. Moreover, BMP, Nodal, and FGF morphogen gradients regulate cell migration independently of cell fate ^51–55^. Thus, it seems that mesodermal cells respond to morphogen cues with precision, providing determinism to the morphogenetic cell behaviours. Simultaneously, they demonstrate plasticity regarding their final fate ^56^. While progenitors are seen giving rise to only one cardiac cell type, they could potentially generate additional cardiac fates when no longer constrain by positional cues. We propose that achieving a delicate balance between determinism and plasticity is essential to ensure robust morphogenesis. This balance enables cells to follow specific developmental pathways while also maintaining the flexibility needed to adapt to changing external cues.

Together, our live-imaging analysis of migration and cardiac lineages provides evidence that some regulation of directionality of cell movements and fate allocation may exist early within the mesoderm. The findings have broader implications for our understanding of organogenesis since they address how initial differences between progenitors and signalling cues may ultimately affect the fate and movements of cells.

## Material and Methods

### Experimental model and subject details

All animal procedures were performed in accordance with the Animal (Scientific Procedures) Act 1986 under the UK Home Office project licenses PP8527846 (Crick) and PP3483414 (UCL) and PIL IA66C8062.

### Mouse strains

The *T^nEGFP-CreERT^*^2^*^/+^*(MGI:5490031) lines were obtained from Hiroshi Sasaki. The *R26 ^TomatoAi^*^14/*Tomato Ai*14^ (*Gt(ROSA)26Sor^tm1^*^4^*^(CAGtdTomato)Hze^* (MGI:3809524), were obtained from the Jackson Laboratory. The *(no gene)^Tg(BRE:H2B;Turquoise)Jbri^* BMP reporter line was generated by the Briscoe laboratory previously ^7^.

### Generation of the cTnnT-2a-eGFP line

The C -terminal tagging of cardiac troponin cTnnt2 with eGFP was generated in the Genetic Modification Service using a CRISPR-Cas9 strategy. This editing was performed by co-transfection of a Cas9-gRNA vector and a donor vector comprising the T2A self-cleaving peptide and eGFP into B6N 6.0 embryonic stem cells using Lipofectamine 2000. The donor vector contained a 786bp insert of T2A-eGFP with 1kb homology arms either side. The guide sequence used was 5’-TTTCATCTATTTCCAACGCC -3’. Two correctly targeted ESC clones were microinjected into blastocysts which were then transferred into the uterus of pseudo pregnant BRAL (C57BL/6 albino) females. Generated chimeras were crossed to BRAL and F0 chimera offspring were initially screened for the proper integration of T2A - eGFP by Sanger sequencing. The F0 were again crossed to BRAL to confirm germline transmission of the mutation and F1 mice were validated by Sanger sequencing.

### Immunostaining

Embryos were dissected in 1x Phosphate buffered saline (PBS, Invitrogen), fixed for 4 hours in 4% PFA at room temperature, washed with 1x PBS-0.1% Triton (0.1% PBS-T) and permeabilised in 0.5% PBS-T. Embryos were blocked with 1% donkey serum for 1hr, incubated overnight at 4°C with antibodies diluted in 0.1% PBS-T: mouse anti-cTnnT (1:250, Thermo Fischer Scientific Systems, MS295P0). Embryos were washed in 0.1% PBS-T at room-temperature and incubated overnight at 4°C with secondary antibodies coupled to 555 fluorophores (1:200, Molecular Probes). After washing with 0.1% PBS-T, embryos were then incubated overnight at 4°C with DAPI (1:1000). Embryos were washed in 0.1% PBS-T and mounted in Rapiclear. Confocal images were obtained on an inverted Sp5 confocal microscope with a 20X oil objective (for early E7.5 embryos) or a 10X air objective (0.4 NA) (for E12.5 hearts) at a 2048 × 2048 pixels dimension with a z-step of 1 to 5 μm. Embryos were systematically imaged throughout from top to bottom. Images were processed using Fiji software ^57^.

### HCR

Whole-mount RNAscope fluorescent in situ hybridizations (ISH) were conducted using the HCR™ RNA-FISH Technology. Wild-type embryos were fixed in 4% PFA overnight and dehydrated in graded MeOH/PBS-Tween 0.1% washes (25%, 50%, 75%, 2×100%) on ice before storage at −20°C. Embryos were rehydrated through a series of graded MeOH/PBS-Tween washes, followed by digestion with proteinase K (20 mg/mL) for 5 minutes (cardiac crescent stage) or 10 minutes (heart tube stage). Embryos were washed twice in PBS-Tween 0.1% and post-fixed with 4% PFA for 20 minutes at room temperature. Post-fixation, embryos were washed in PBS-Tween (3×5 min) and pre-incubated in a 50/50 hybridization buffer and PBS-Tween mix for 5 minutes. Embryos were then incubated with probe hybridization buffer preheated to 37°C and hybridized overnight at 37°C with specific probes (1 μM). Embryos were washed in preheated probe wash buffer at 37°C (2×5 min, 2×30 min), followed by a 10 minutes wash in a 50/50 probe wash buffer and 5X SSCT at 45°C. After three 5 mininutes washes in 5X SSCT at room temperature, embryos were incubated in 1 mL amplification buffer for 30 minutes at room temperature. Fluorescent hairpin solutions (3 μM each) were snap-cooled and prepared in amplification buffer, and embryos were incubated overnight in the dark at room temperature. Excess hairpins were removed by washing with 5X SSCT (2×5 min, 2×30 min, 1×5 min) and stored at 4°C protected from light. Embryos were stained with DAPI overnight, washed (3×10 min, 5X SSCT), and mounted on slides using Rapiclear solution. Confocal microscopy was performed using a 20x oil immersion lens, with a step size of 1 μm for confocal imaging (Leica Sp5). The following probes were used: B4-Nr2f2 NM_183261.3; B5-HCN4 NM_001081192.2 and B2-cTnnT2 NM_001276345.2.

### Lineage tracing

To synchronize estrous cycles, females were exposed to male-soiled bedding for 3 days prior to mating. On the fourth day, mating was conducted for 3 hours (from 8 AM to 11 AM), and the presence of vaginal plugs was checked at 11 AM, designating the embryonic stage as E0. Tamoxifen (T5648, Sigma) was administered via oral gavage at a dose of 0.02 mg/g body weight, dissolved in corn oil, at specific developmental stages.

### Embryo harvesting and culture

Embryos were collected at E6.5, with morning dissections (∼8 AM) for embryo imaged in Movie 1 and afternoon dissections (5-7 PM) for embryos imaged in Movies 2-5. The embryo dissected in Movie 1 was at an earlier stage than the embryo in Movie 5, yet both were dissected in the afternoon at similar times. This variation is due to the high degree of variability in embryonic stages within each litter and between different litters, as previously documented^7^. Harvesting was performed in DMEM (D5921, Sigma-Aldrich) containing 10% FBS (A5256701, Thermo Fisher), 25 mM HEPES-NaOH (pH 7.2), penicillin, and streptomycin. Dissections were conducted under a stereoscope equipped with a Tokai Hit thermoplate set at 37°C and completed within 5 minutes to preserve developmental potential. Embryos were then transferred to a culture medium consisting of 75% freshly prepared rat serum (filtered through a 0.2 μm filter) and 25% DMEM (supplemented with 1 mg/mL D-glucose and pyruvate, without phenol red and L-glutamine, D5921, Sigma-Aldrich), further enriched with GlutaMAX, 100 U/mL penicillin, 100 μg/mL streptomycin, and 11 mM HEPES. Rat serum was prepared according to established protocols, heat-inactivated at 56°C for 30 min, and stored at −80°C until use. The culture medium was equilibrated in 5% O2, 5% CO2, and 90% N2 atmosphere and pre-warmed to 37°C before embryo transfer.

### Embryo positioning and multiphoton imaging

Custom plastic holders were fabricated with varying hole diameters (0.3–0.5 mm) to secure the ectoplacental cone, following the method described by Nonaka et al. ^58^. Embryos were mounted with the anterior side facing up and overlaid with mineral oil (M8410, Sigma-Aldrich) to prevent medium evaporation. Before imaging, embryos were pre-cultured for 2 hours in the microscopy setup. Time-lapse imaging was performed using a multiphoton Olympus FVMPE-RS microscope equipped with a 5% CO2 incubator and a heated chamber maintained at 37°C. A 20x/1.00 W dipping objective with a 2-mm working distance was used for imaging. A SpectraPhysics MaiTai DeepSee pulsed laser set at 880 nm was used for single-channel two-photon imaging. Image acquisition settings included 250 mW output power, 7 μs pixel dwell time, and 610 × 610 μm image dimension (1,024 × 1,024 pixels). The z-step was set to 6 μm.

### Light sheet microscopy

For all light sheet acquisitions, we imaged *TnEGFP-CreERT2/+*; *R26 Tomato Ai14*/ *Tomato Ai14*, *cTnnT-2a-GFP+/−* embryos. Tamoxifen (T5648, Sigma) was administered via oral gavage at a dose of 0.02 mg/g body weight dissolved in corn oil at specified embryonic stages, and embryos were harvested at least 12 hours post-gavage. Prior to imaging, embryos were pre-cultured for 2 hours in the microscopy setup. To maintain positional stability during imaging, the ectoplacental cone was partially embedded in growth factor-reduced, phenol red-free Matrigel (356231, Corning) diluted 1:1 with culture medium within a dedicated open-top FEP sample chamber containing four wells. One embryo was mounted per well, accommodating up to four embryos per experiment. Imaging was conducted using the Viventis LS1 light sheet microscope with a single-view, dual illumination configuration. Detection was performed using a Nikon 25x NA 1.1 water immersion objective (final magnification: 18.7x), yielding an 800 × 800 μm field of view. Sequential illumination was applied with 488 nm and 561 nm lasers. Image acquisition was performed every 2 minutes with z-stacks spanning 500 μm. The voxel size was 2 μm (z) × 0.347 μm (x, y), and the light sheet thickness was 3.3 μm. Exposure times for GFP and tdTomato channels were set to 50 ms and 50 to 100 ms, respectively. Embryos were maintained at 37°C in an 8% CO2 humidified environment throughout the imaging session.

### Cell tracking

The original images were converted into HDF5/XML file formats and registered using BigStitcher [14], following previously established methodologies [6]. Processed images were then imported into the MaMuT Fiji plugin for manual cell tracking [5]. The MaMuT viewer allowed visualization of tracked cells with adjustable brightness, scale, and rotation settings. Cell identification was initiated at the first time point at which the cell’s tdTomato fluorophore starts to be detectable in a movie, with manual tracking performed every five time-points, except during mitosis, where tracking was conducted at every time point to ensure accurate measurements of daughter cell separation. The TrackScheme lineage browser was used to visualize reconstructed cell lineage trees, representing cells as nodes interconnected by edges, and cell divisions as split branches. For subsequent analysis, tracked cells’ positions, tracks, division times, and GFP intensities were exported in CSV format.

### Threshold analysis

GFP values were normalised based on the highest GFP value within each movie’s tracking data. To discern GFP-positive cells from the background, we implemented the following approach: GFP background intensities were measured at regular intervals throughout each movie and linear interpolation was applied between time points to establish a background value for every time-point. The threshold for distinguishing GFP-positive and GFP-negative cells was determined by using a multiple of the mean background. The threshold was set to the lowest multiplier, ensuring that all endocardial cells in each movie were classified as GFP-negative.

### Temporal alignment of the time-lapse movies

The movies were temporally aligned based on the differentiation timing of LV/AVC and inflow myocytes. Differentiation was defined as the point when the cTnnT-2a-eGFP signal intensity surpassed a predefined threshold. Timepoint 0 in Movie 2 was used as the reference (T0). The time in the other movies was adjusted relative to this reference by adding 74 minutes (Movie 1), 239 minutes (Movie 3), 373 minutes (Movie 4), and 626 minutes (Movie 5) to synchronize their differentiation timings. All lineage trees were reconstructed using these adjusted time points. Birth date is the time point at which the cell’s tdTomato fluorophore starts to be detectable in a movie.

### Lineage analysis

Each reconstructed lineage tree starts with a mother cell. The mother divides, giving rise to the two daughter cells D1 and D2. Then D1 divides giving rise to daughters D11 and D12. D11 divides giving rise to daughters D111 and D112. Lineage trees were generated in R version 4.3.2 using the ‘ggtree’ version 3.10.0 package with Newick format and imported into R through the ‘read.tree’ function from the ‘ape’ package version 5.7.1. The branch lengths of the trees are proportional to the cell cycle length. The trees were coloured using the ‘scale_colour_gradient’ function in the ‘ggtree’ package, which imparts a colour gradient based on the normalised GFP intensities, speeds or tortuosity of the cells or paintsubtree function in phytools version 2.03 package in R version 4.3.2 ^59, 60, 61^.

### Analysis of myocardial clone dispersion

To analyze the spread of myocardial clones along the anterior-posterior (AP) and dorsal-ventral (DV) axes, we first empirically established these axes in the heart tube. We then identified all cTnnT-2a-eGFP positive cells for each clone and mapped their coordinates onto the AP and DV axes. The spread of each clone was quantified as the proportion of its range along these axes. To reduce bias from short-range movements immediately following cell division, we included only cells that were at least 4 hours into their cell cycle in the analysis.

### Quantification of sister cell contact duration

We quantified the proportion of time that sister cells remained in contact following cell division. To determine the contact period, we measured the distances between 57 pairs of cells from Movies 1 and 2, which were visually confirmed to be in contact. The mean contact distance was 13 μm. Distances between sister cells were then measured at each time point, and the duration for which the distance remained below 13 μm was recorded. The cell contact duration was expressed as a proportion of this contact period relative to the total cell cycle length. Only sister cells from generations 1 (D1, D2) and 2 (D11, D12, D21, D22) were included in the analysis. The study was restricted to the early migration period, within the first 16 hours of the aligned movies, corresponding to the timeframe when the first myocardial cell becomes cTnnT-2a-GFP+. Sister cells with a cell cycle duration of less than 4 hours were excluded.

### Homotypic and heterotypic distances calculation

Trajectories were reconstructed in 3D using the matplotlib version 3.7.2 and plotly version 5.18.0 packages in Python version 3.11.53. Euclidean distances were computed to quantify the spatial relationships among cells belonging to the same family tree, categorizing them as ‘homotypic distances’ for cells of the same fate and ‘heterotypic distances’ for cells of different fates. Progenitors’ coordinates were sampled at two distinct time points: at the end of migration and during the stages when the heart tube forms. We only considered branches lasting at least 4 hours into the cell cycle to allow sufficient cell migration. The migration period for early proximal mesoderm cells, which include the LV/AVC, ExEm, endocardial, and pericardial cells, was defined as the time from the beginning of the movie until the completion of cell migration, marked by the mean differentiation time of the LV/AVC region. The mean differentiation times for LV/AVC cells were 20.7 hours for Movie 1, 20.8 hours for Movie 2, 17.3 hours for Movie 3, 16.3 hours for Movie 4, and 11.5 hours for Movie 5. The subsequent heart tube morphogenesis period extended from the end of the migration phase until the end of the movie, corresponding to the stages of heart tube formation. For atrial cells, the migration period concluded at the mean atrial differentiation time. The mean differentiation times for inflow cells were 29.1 hours for Movie 1, 26.8 hours for Movie 3, 23.3 hours for Movie 4, and 19.8 hours for Movie 5. No inflow cells were tracked in Movie 2.

To analyze the evolution of homotypic and heterotypic distances over time, we focused on daughters’ coordinates at three distinct time points: T0, T1, and T2. T0 was 20 minutes after the mother’s initial cell division, T1 was the last time point before the daughters’ subsequent cell division, and T2 was the final time point at which all granddaughter cells existed before the end of migration (as described above). If no third generation was present, we only considered branches lasting at least 4 hours into the cell cycle to allow sufficient cell migration.

### Speed calculation

To compare the motility of different cell types, we calculated their speed using the CelltrackR package version 1.1.0 in R version 4.3.2 ^62^. Speed was determined by dividing the total track length of each cell by its duration, resulting in a mean speed expressed in μm/minute. To evaluate changes in speed over time, the aligned movies were divided into 5-hour intervals, and the mean speed of cells in each interval was calculated. Cells with a cell cycle length of less than 40 minutes within any interval were excluded from the analysis.

### Cell tortuosity calculation during heart tube formation

Tortuosity quantifies the extent to which a cell’s path deviates from a straight line. It was calculated using the ratio of the shortest straight-line distance between two points to the total path distance covered by the cell:

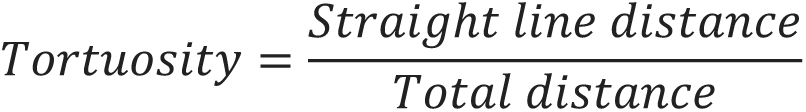

A tortuosity value closer to one indicates a straighter migratory path, while values approaching zero signify a more curved trajectory. Local tortuosity was calculated at 50-minute intervals and then averaged over the cell cycle duration computed for each cell. Cells with a cell cycle length of less than 3 hours were excluded from the analysis.

### Dynamic time warping

Migratory trajectories were analyzed by comparing the paths of sister cells from the point immediately after their mother cell division to the final time point before the first sister cell divided again. Trajectory similarity was assessed using dynamic time warping (DTW) distance, calculated using the dtw-python package (version 1.3.0). DTW was chosen over Euclidean distance because it assign short distsances to spatially similar but temporally asynchronous trajectories. The analysis used the ‘symmetricP1’ step pattern, a slope-constrained step pattern ^63,64^. DTW values were calculated separately for sister cells with the same fate (unipotent) and different fates (bipotent) across the first and second generations (D1, D2; D11, D12, D21, D22). Distinct analyses were performed for progenitors contributing exclusively to mesoderm (Uni_meso and Bi_meso) and progenitors contributing to at least one extraembryonic mesoderm (ExEm) daughter (Uni_ExEm and Bi_ExEm). To measure trajectory similarity during the migration period, DTW values were calculated up until the end of migration. To determine whether DTW values differed significantly between unipotent and bipotent sister cells over time, mean DTW values for the first 40 steps were plotted, and statistical comparisons were made using the Mann-Whitney test.

### Permutation test

We generated 100,000 permutations by pooling all log DTW distances and randomly assigning them to either unipotent or bipotent conditions. For each permutation, we calculated the difference in mean log DTW values between sister cells with shared fates and those with distinct fates. This iterative process created a null distribution, allowing us to test whether unipotent progenitors produce sisters with more similar trajectories than bipotent progenitors (Figure 6 Supplementary Figure H-K).

## Acknowledgments

We thank Dr Martin Dominguez for his initial help with BigStitcher. We thank Dr Rosie Marshall and Prof. Andrew Copp for their help with rat serum preparation. We also thank the Light Microscopy facilities and Biological Research Facilities at the Francis Crick Institute and UCL for their help with imaging and transgenic colonies.

## Author Contribution

S.A. Conceptualization, Formal analysis, Investigation, Visualization, Writing – review & editing, P.A.E. Methodology, Writing – review & editing, A.C. Methodology, S.V.B Methodology, I.N. Investigation, J.A.D. Methodology, Supervision, Writing – review & editing, J.B. Funding acquisition, Writing – review & editing, K.I. Funding acquisition, Project Administration, Conceptualization, Formal analysis, Investigation, Visualization, Supervision, Writing – original draft, Writing – review & editing.

## Data Availability

All relevant data source and codes used are available at database: https://github.com/sabukar22/Supplementary-Data

---

**Figure 2 – Supplementary Figure 1.**
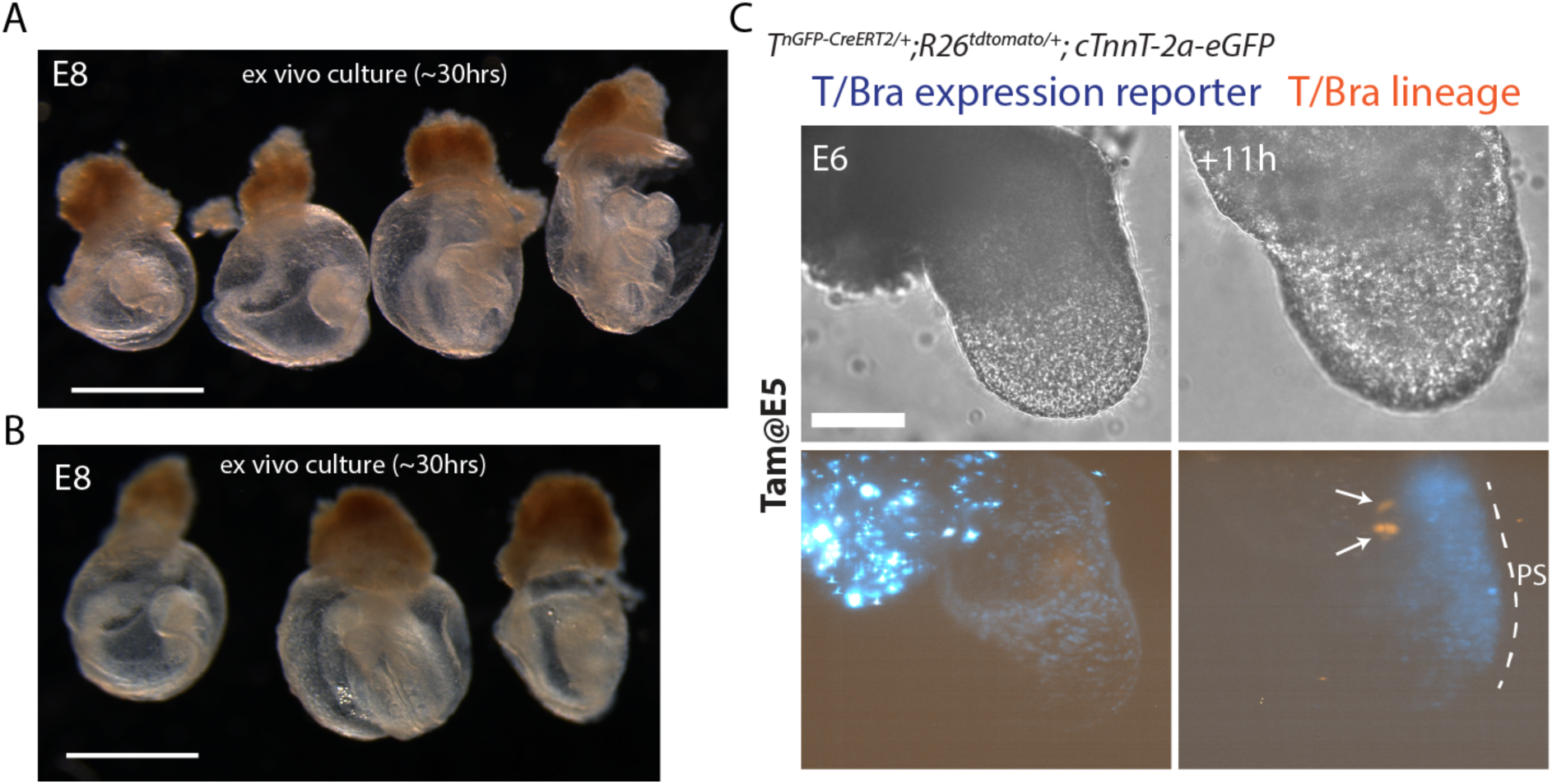
Embryo Culture and Tamoxifen Administration. (**A-B**) Two independent embryo culture for 30 hours. Scale bar: 1 mm (**C**) Time-lapse of *T^nGPF-CreERT^*^2^*^/+^;R26R^tdTomato/+^; cTnnT-2a-eGFP* embryos after tamoxifen administration (0.02 mg/body weight) at E5, followed by 11 hours of culture from E6.0 without tamoxifen. Arrows indicate tdTomato-positive cells in the extra-embryonic mesoderm. Scale bar: 100 μm. bw: body weight.

**Figure 2 – Supplementary Figure 2.**
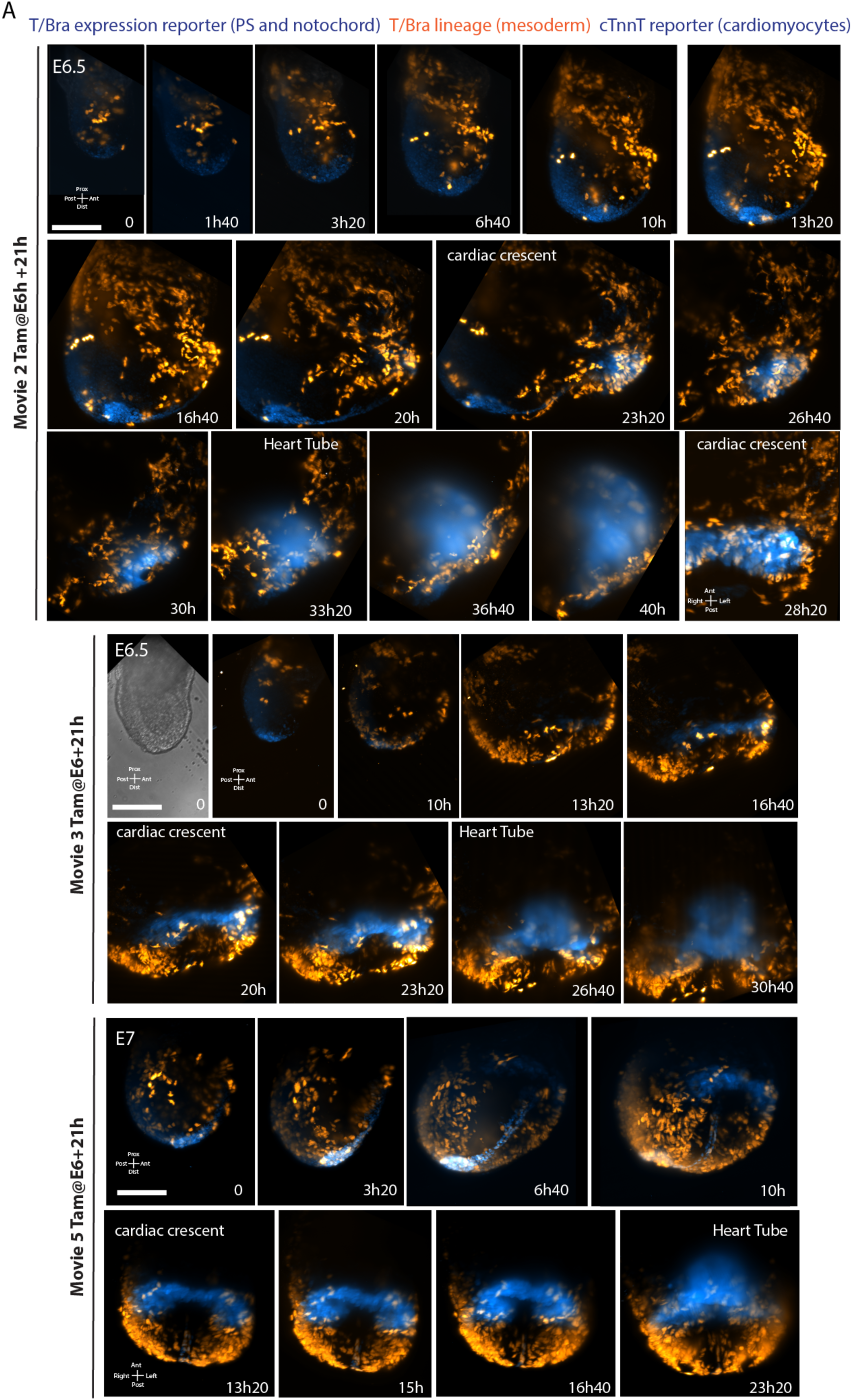
Long-term live-imaging from gastrulation to heart tube formation. (**A**) Time-lapse sequences from *T^nGPF-CreERT^*^2^*^/+^;R26R^tdTomato/+^; cTnnT-2a-eGFP* embryos after tamoxifen administration (0.02 mg/body weight) at the indicated stage. LV/AVC indicates the left ventricle and atrioventricular canal. Images from A-movie 2 at 28:20 and movie 5 from 13:20 to 23:20 were computationally rotated in BigStitcher to show the ventral side. Scale bar: 100 μm. Ant: Anterior; Pos: Posterior.

**Figure 3 – Supplementary Figure 1.**
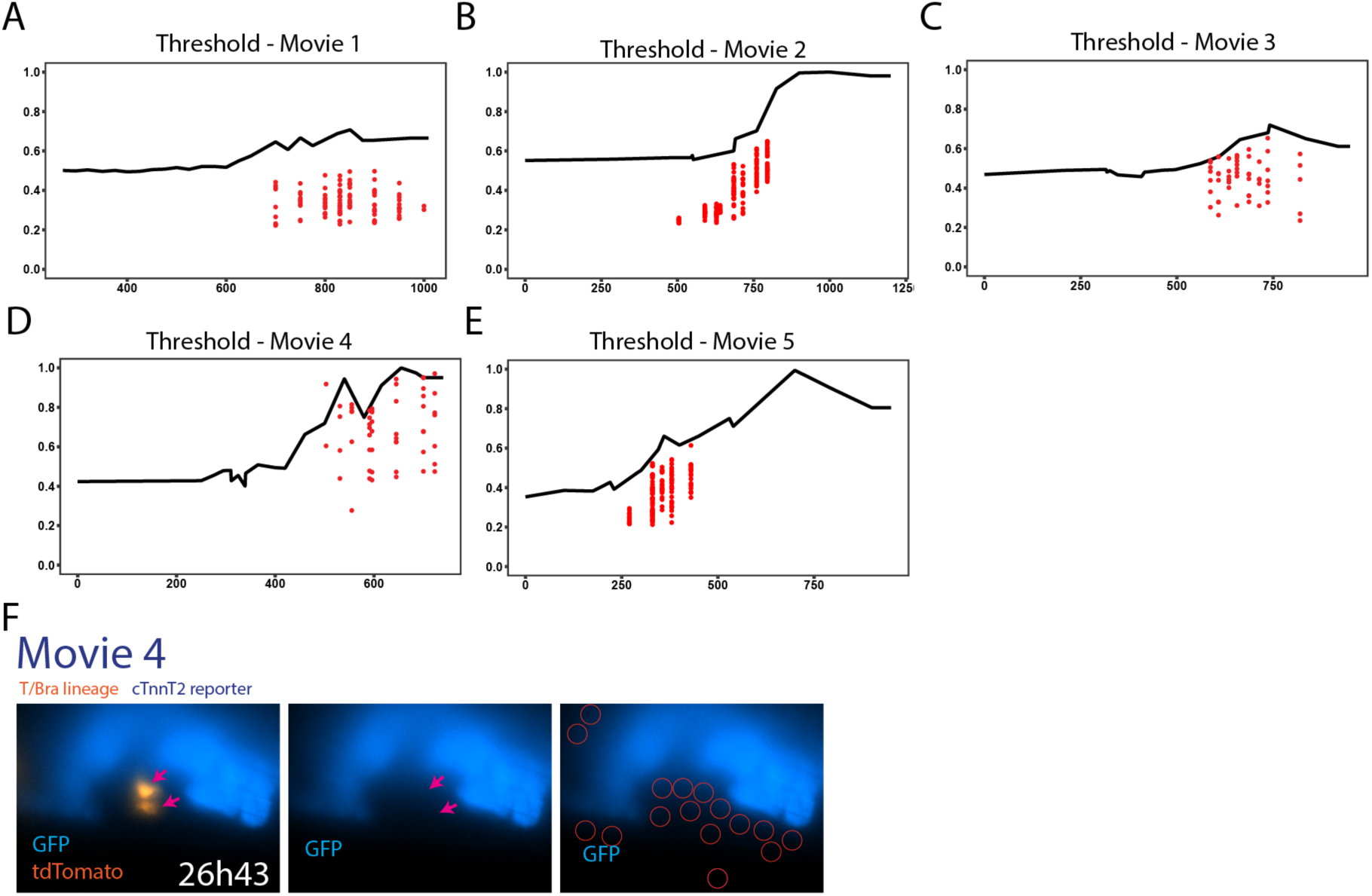
Threshold Analysis. (**A-E**) Black lines denote the GFP intensity threshold separating GFP-positive cells from the background. Red points show GFP intensity values of endocardial cells in each movie. (**F**) Red circles indicate areas used to measure background GFP intensity.

**Figure 4 – Supplementary Figure 1.**
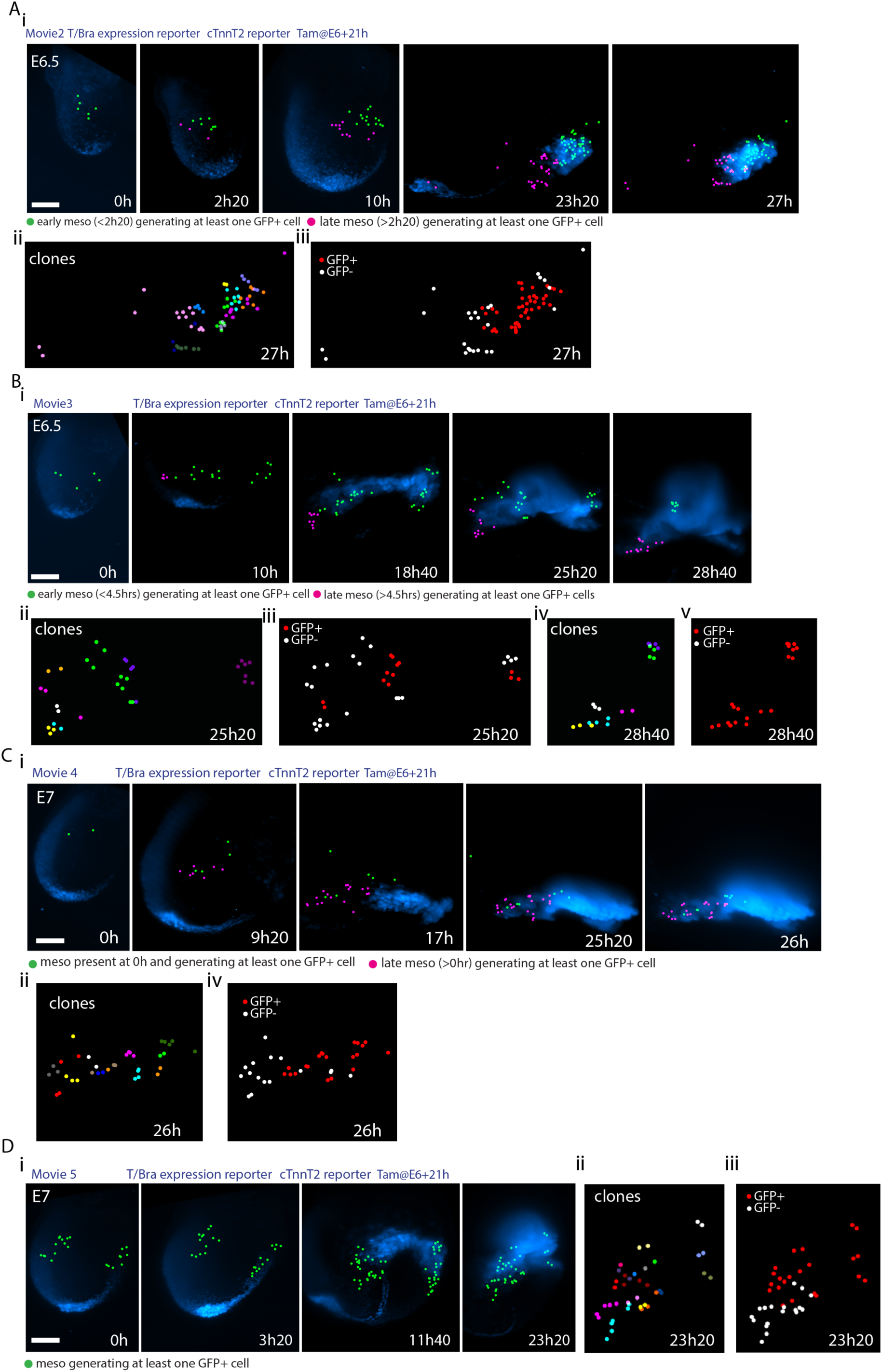
Independent progenitors contribute to the LV/AVC and Atria. (**A-D**) Only progenitors contributing at least one cTnnT-2a-GFP+ cell are displayed. (i) Fate map showing early (green) and late (magenta) mesoderm contributions to heart tube regions (early/late classification is arbitrary). (ii) Each color represents a unique clone. (iii) cTnnT-2a-GFP+ cells in red. Scale bar: 100 μm.

**Figure 5 – Supplementary Figure 1.**
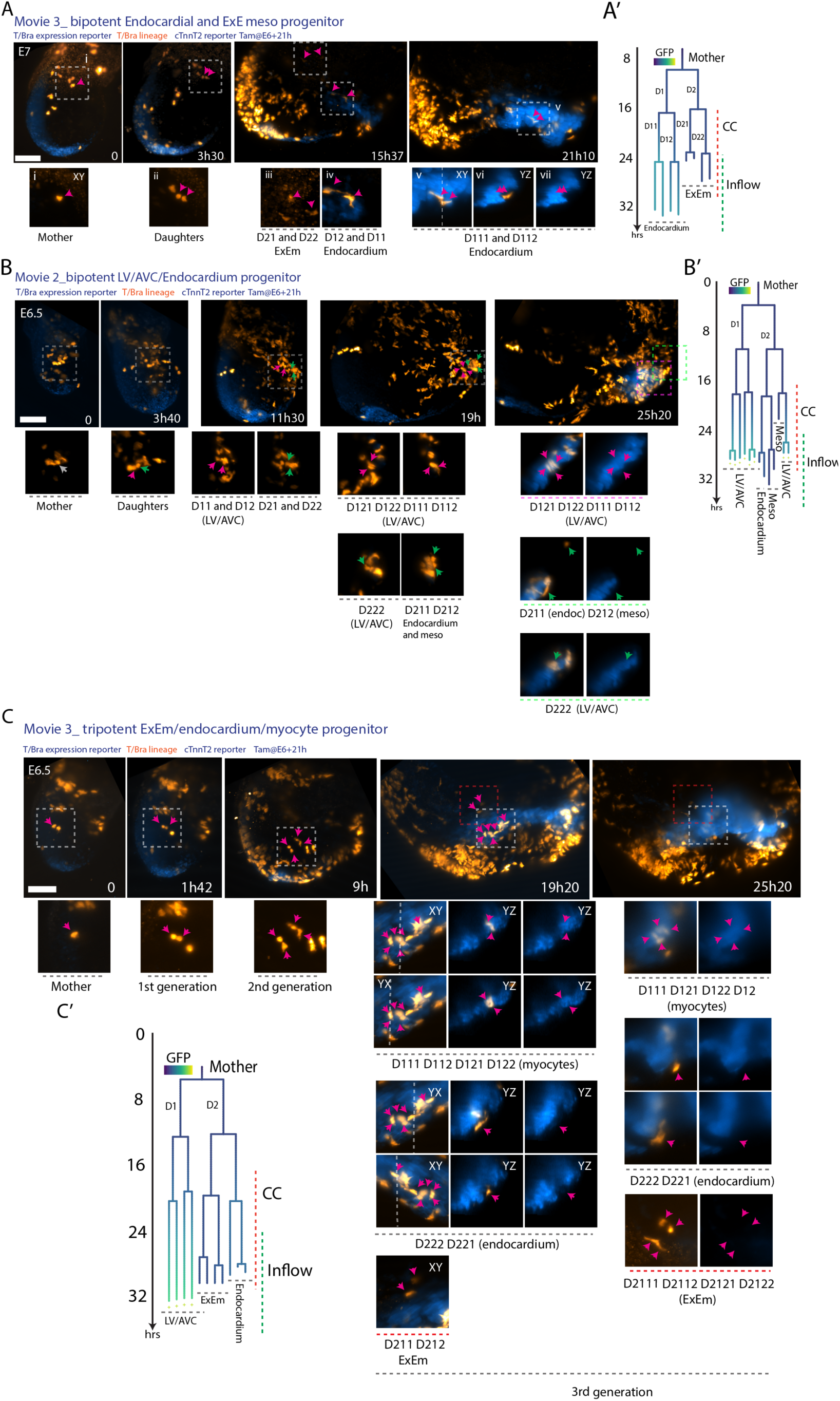
Examples of Bipotent and Tripotent mesodermal progenitors. (A-C) Time-lapse images of *T^nGPF-CreERT^*^2^*^/+^;R26R^tdTomato/+^; cTnnT-2a-eGFP* embryos showing bipotent (A-B) and tripotent (C) progenitors. Corresponding lineage trees, colored by normalized GFP intensity, are shown in (A’-C’), with arrows indicating the cells in (A-C).

**Figure 5 – Supplementary Figure 2.**
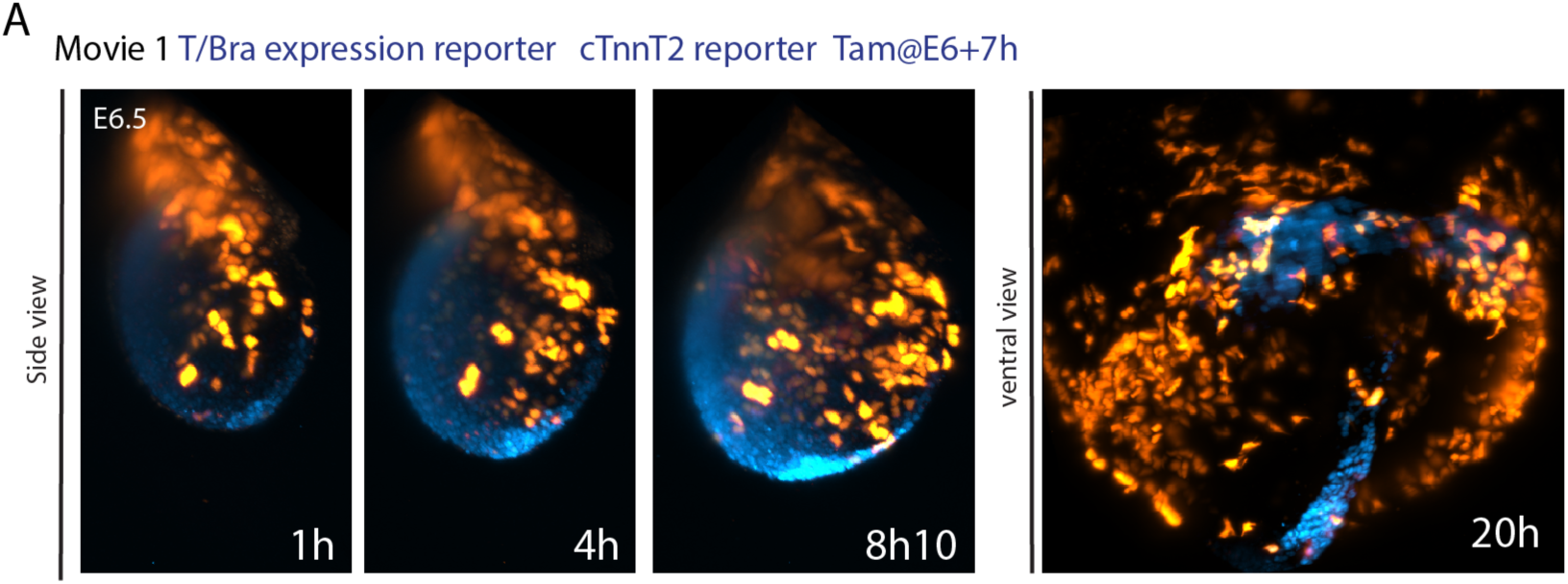
Movie 1. **A.** Images corresponding to Figure 5A, with additional labeling to show T/Bra lineage-positive cells marked by tdTomato fluorescence.

**Figure 6 - Supplementary Figure 1.**
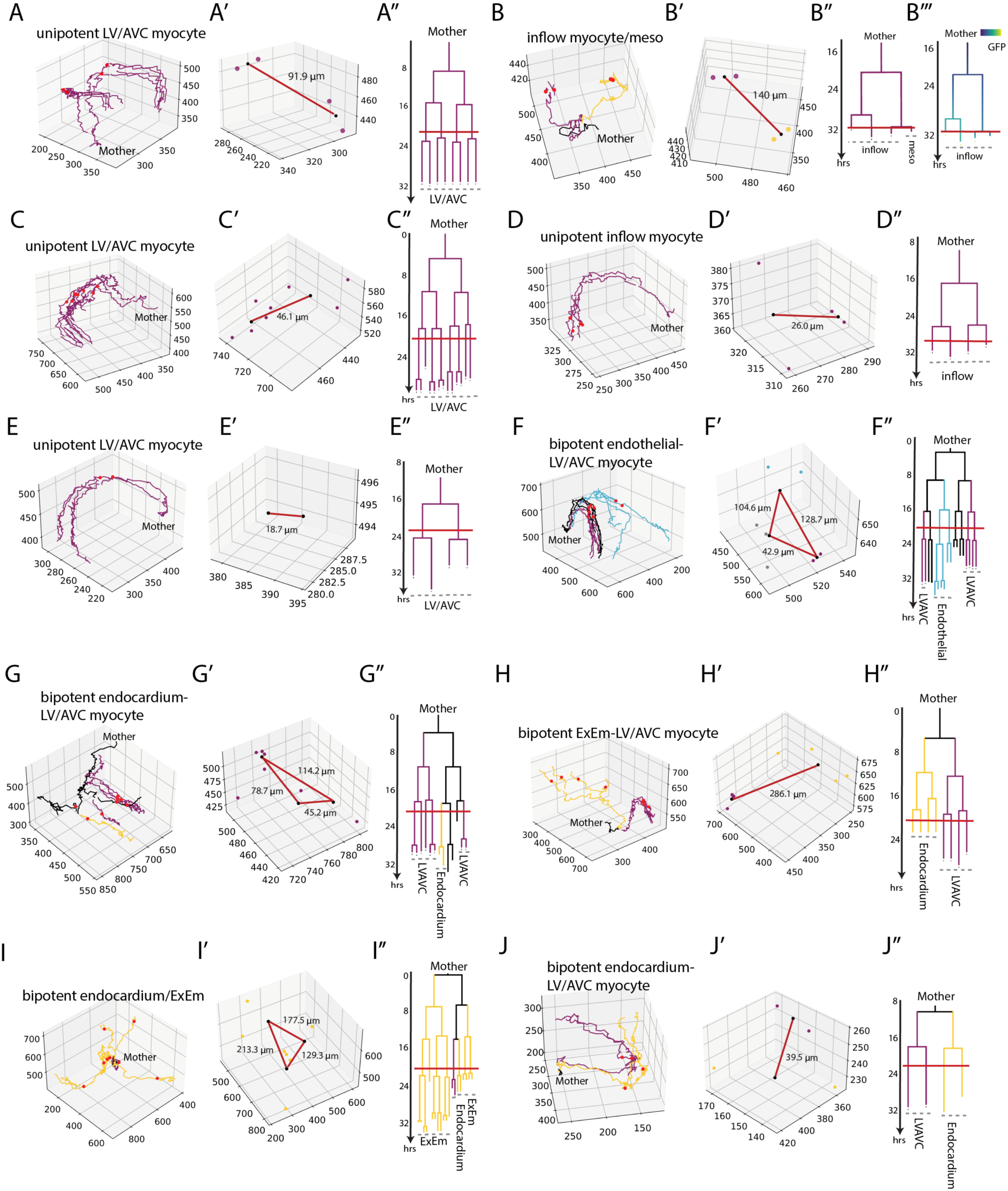
Mesodermal migration paths. Examples of trajectories (A-K) and corresponding midpoint distances (A-K) and lineage trees (A”-K”). Red lines mark sampling points for midpoint analysis. LV/AVC: left ventricle and atrioventricular canal; ExEm: Extra embryonic mesoderm.

**Figure 6 - Supplementary Figure 2.**
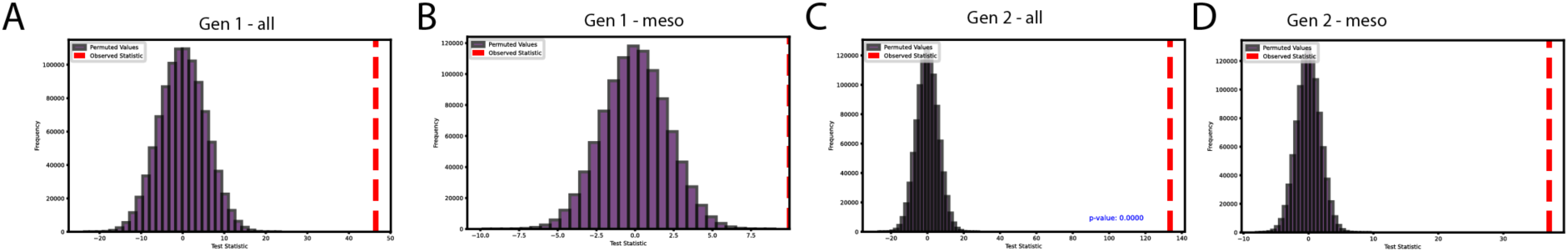
Statistical tests (A-D) Histograms showing the distribution of permuted tests based on the log mean difference between randomly shuffled unipotent and bipotent DTW distances for: all D1 and D2 cells (A), D1 and D2 cells excluding ExEm progenitors (B), all D11, D12, D21, and D22 cells (C), and D11, D12, D21, and D22 cells excluding ExEm progenitors (D). Data are binned at 30 µm intervals, with the red dashed line indicating the observed log mean difference.

**Figure 6 - Supplementary Figure 3.**
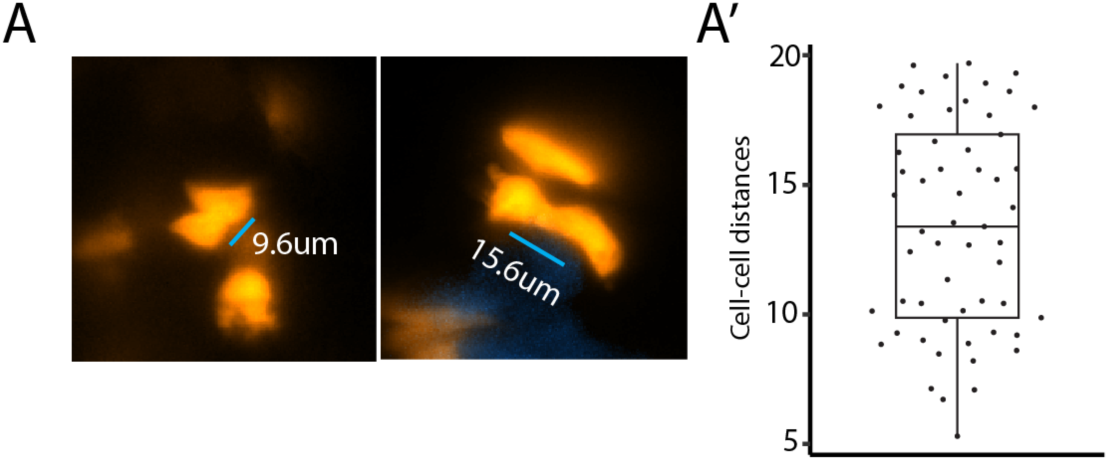
**Cell-cell contact measruements. (**A) Example of cell-cell distance measurement between cells that are in contact. (A’) Mean cell-cell distances.

**Figure 7 - Supplementary Figure 1.**
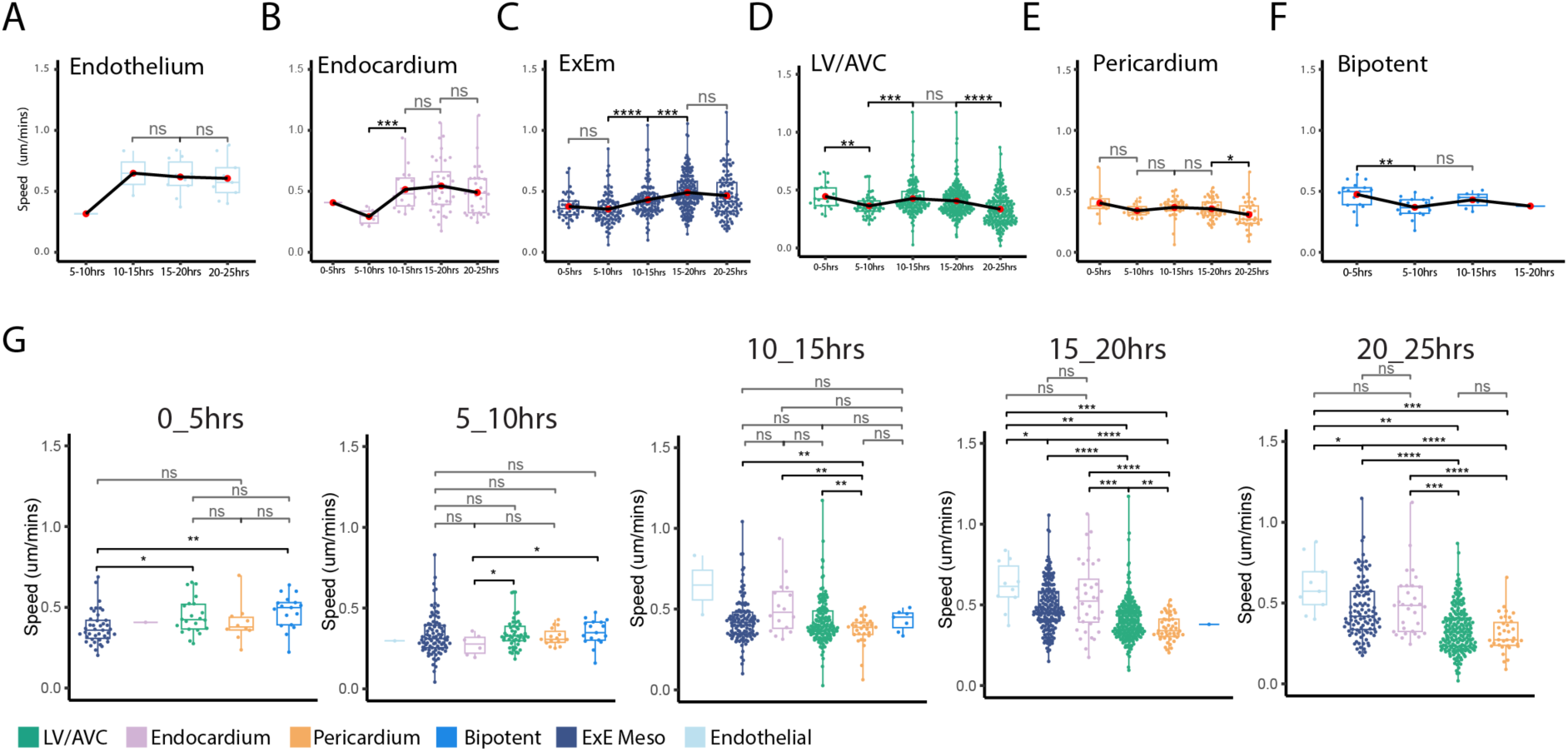
Cell speed analysis. (**A-E**) Cell speeds per cell fate were calculated across 5-hour time-periods. Movies were temporally aligned as shown in Figure 4B. See also Figure 6H. All statistical analyses were performed using the Mann-Whitney U test. LV/AVC: left ventricle and atrioventricular canal; ExEm: Extraembryonic mesoderm.

**Figure 7 - Supplementary Figure 2.**
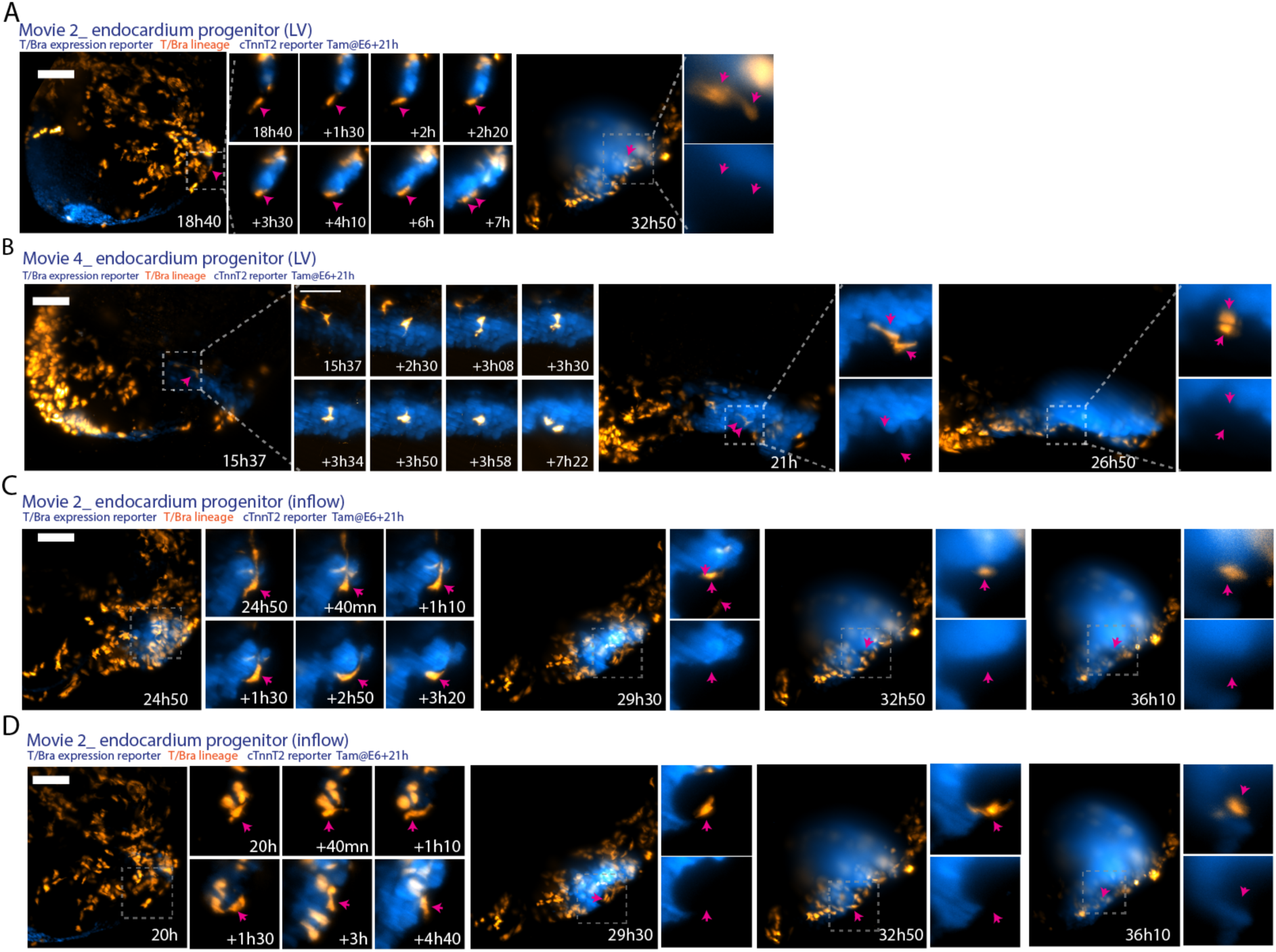
Endocardial cell behaviour. (**A-D**) Image sequences from time-lapse videos of *T^nGPF-CreERT^*^2^*^/+^;R26R^tdTomato/+^; cTnnT-2a-eGFP* embryos showing endocardial progenitors in the LV (A-B) and in the inflows (C-D). LV: left ventricle.

**Figure 7 - Supplementary Figure 3.**
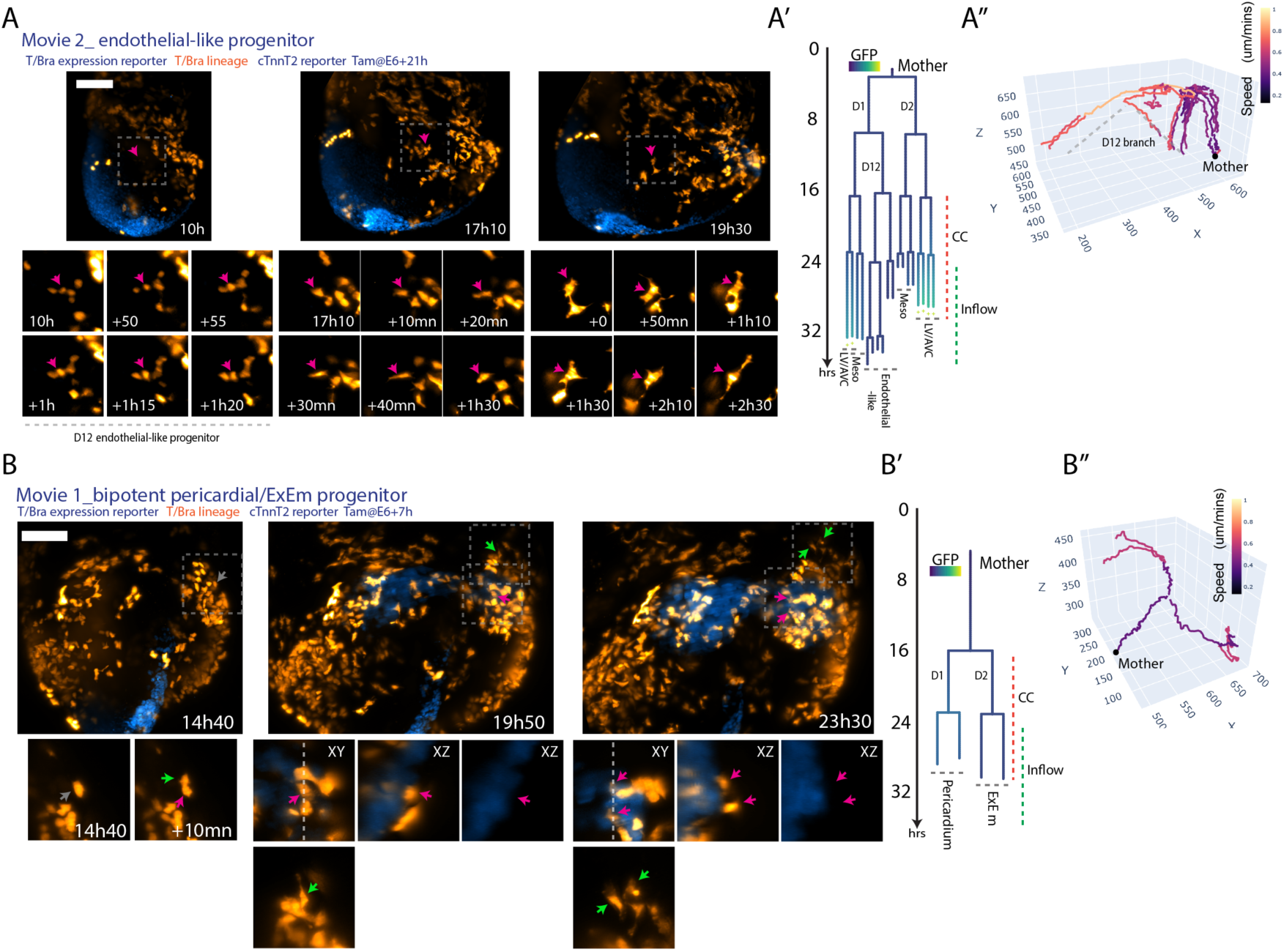
Mesodermal cell behaviours (A-B) Time-lapse images of *T^nGPF-CreERT^*^2^*^/+^;R26R^tdTomato/+^; cTnnT-2a-eGFP* embryos showing an endothelial-like progenitor (A) and a bipotent pericardial and ExEm progenitor (B). **(A)** Endothelial-like progenitor with progressive shape changes; lineage tree colored by normalized GFP intensity shown in (A’), with pink arrow indicating cell in branch D12. 3D plots (A’’) color-coded by cell speed. **(B)** Bipotent pericardial and ExEm progenitor; lineage tree with normalized GFP intensity shown in (B’), with arrows indicating cells. 3D plot (B’’) color-coded by cell speed. ExEm, extra-embryonic mesoderm; CC, cardiac crescent; Meso, mesoderm; LV/AVC, left ventricle-atrioventricular canal. Scale bar: 100 μm.

**Video 1-2:** Live-imaging from gastrulation to heart tube stage and cell tracking. Related to Figure 2B and Figure 2-supplementary Figure 2A.

**Video 3:** Example of a myocyte progenitor visualised along the xy and yz axis. Scale bar: 50 μm.

**Video 4:** Example of an endocardial progenitor visualised along the xy and yz axis. Scale bar: 50 μm.

**Video 5:** Example of a pericardial progenitor visualised along the xy and yz axis. Scale bar: 50 μm.

**Video 6**: 3d reconstruction of regional landmarks traced from cardiac crescent to heart tube stage. Related to Figure 2D.

**Video 7**: 3d reconstruction of clones traced from gastrulation to heart tube stage. Related to Figure 4G.

**Video 8**: Fate mapping of the early mesoderm. Related to Figure 5A.

